# Integrated genomic and functional analyses of human skin-associated *Staphylococcus* reveals extensive inter- and intra-species diversity

**DOI:** 10.1101/2023.06.22.546190

**Authors:** Payal Joglekar, Sean Conlan, Shih-Queen Lee-Lin, Clay Deming, Sara Saheb Kashaf, NISC Comparative Sequencing Program, Heidi H. Kong, Julia A. Segre

## Abstract

Human skin is stably colonized by a distinct microbiota that functions together with epidermal cells to maintain a protective physical barrier. *Staphylococcus*, a prominent genus of the skin microbiota, participates in colonization resistance, tissue repair, and host immune regulation in strain specific manners. To unlock the potential of engineering skin microbial communities, we aim to fully characterize the functional diversity of this genus within the context of the skin environment. We conducted metagenome and pan-genome analyses of isolates obtained from distinct body sites of healthy volunteers, providing a detailed biogeographic depiction of staphylococcal species that colonize our skin. *S. epidermidis*, *S. capitis,* and *S. hominis* were the most abundant species present in all volunteers and were detected at all body sites. Pan-genome analysis of these three species revealed that the genus-core was dominated by central metabolism genes. Species-specific core genes were enriched in host colonization functions. The majority (∼68%) of genes were detected only in a fraction of isolate genomes, underscoring the immense strain-specific gene diversity. Conspecific genomes grouped into phylogenetic clades, exhibiting body site preference. Each clade was enriched for distinct gene-sets that are potentially involved in site tropism. Finally, we conducted gene expression studies of select isolates showing variable growth phenotypes in skin-like medium. *In vitro* expression revealed extensive intra- and inter-species gene expression variation, substantially expanding the functional diversification within each species. Our study provides an important resource for future ecological and translational studies to examine the role of shared and strain-specific staphylococcal genes within the skin environment.

**SIGNIFICANCE:** The bacterial genus *Staphylococcus* is a prominent member of the human skin microbiome, performing important and diverse functions such as tuning immunity, driving tissue repair, and preventing pathogen colonization. Each of these functions is carried out by a subset of staphylococcal strains, displaying differences in gene content and regulation. Delineating the genomic and functional diversity of *Staphylococcus* will enable researchers to unlock the potential of engineering skin communities to promote health. Here, we present a comprehensive multi-omics analysis to characterize the inter- and intra-species diversity present in human skin-associated staphylococci. Our study is the first to conduct a detailed pan-genome comparison between prominent skin staphylococcal species giving a valuable insight into gene sharing and provides an important resource.

## INTRODUCTION

The human epidermis and associated appendages including hair, nails, and glands (sebaceous, apocrine, eccrine) form a physical and immunologic barrier between our body and the environment, protecting us against pathogens, injury, and unregulated water loss (1). The human skin microbiota, which stably colonize this epidermal interface, play an integral role in skin barrier function via microbe-microbe and host-microbe interactions (2–4). While early studies focused on the pathogenicity of skin microbes, further studies have explored the myriad ways in which microbial strains serve beneficial roles, including providing colonization resistance, wound healing, and tuning immune responses (5). There’s a growing recognition that microbial functions are often strain specific, demonstrating a need to understand the full genetic diversity of the commensal microbiota. For example, strains of *Staphylococcus epidermidis* that produce the serine protease Esp, were shown to limit *S. aureus* colonization by selectively degrading proteins essential for biofilm formation (6). Many staphylococcal strains produce diverse antimicrobial peptides such as lugdunin and phenol-soluble modulins that restricted pathogen growth (7–10). In addition, colonizing *S. epidermidis* strains were shown to directly interact with the host by secreting enzymes that increased the level of skin lipids or ceramides, preventing excessive water loss (11). *S. epidermidis* strains shape and regulate skin immune function through coordinated action of resident immune cells in a highly dynamic fashion (12, 13). Intriguing new studies genetically engineered *S. epidermidis* strains to express antigens which elicit *de novo* immune responses, even demonstrating targeted activity against neo-antigens expressed by melanoma cells (14). Together, these studies focused on *S. epidermidis* and related non-coagulase staphylococci (15) collectively demonstrate the diverse functions encoded by these species to maintain skin health and the future potential to engineer skin communities to restore health.

Despite their strain-dependent contribution to host health, current knowledge about healthy skin-associated staphylococcal genomic diversity is limited. Previous sequencing studies based on 16S rRNA gene amplicon (V1-V3) and shotgun metagenomics have revealed that *Staphylococcus* is one of the most prominent bacterial genera, along with *Cutibacterium* and *Corynebacterium* that persistently colonizes the human skin (2, 16). However, a detailed species-level picture of staphylococci and their distribution across different body sites is currently lacking. Further, building upon a previous pan-genome analysis of commensal *S. epidermidis* isolates (17), strain-level genetic diversity in other healthy skin-associated staphylococci remains unexplored. A pan-genome analysis captures the entire genetic diversity in a given taxon by categorizing genes into core and accessory (18), thus identifying lineage-specific genes under strong selection versus strain-specific genes that encode niche specifying functions. In addition to genetically encoded diversity, studies in other host associated species such as *E. coli* have demonstrated the role of differential gene expression in species diversification (19), thus highlighting the need to study gene expression to further comprehend functional diversity.

In our current work, we seek to study skin-associated staphylococcal diversity from healthy volunteers using metagenomic, pan-genomic, and gene expression analyses to gauge its functional potential. Based on our previous work which established that the V1-V3 based 16S rRNA analysis allows species-level discrimination of staphylococci (20), our current work aimed to characterize the distribution of staphylococcal species across 14 skin sites from 22 healthy volunteers. We show that the skin is colonized by multiple species but predominated by *S. epidermidis*, *S. hominis*, and *S. capitis*, for which human skin serves as the primary habitat (21). We focused on these three predominant species and sequenced 126 non-clonal isolate genomes cultured from 18 skin sites of 14 healthy volunteers, with an aim to understand species and genus level pan-genome diversity. ∼21% genes are core across all species, along with a substantial portion (∼11%) of genes being core in select species, indicating lineage-restricted selection to retain these genes. Most of the genes within the pan-genomes are not core but are variably shared, showing tremendous strain-level diversity as previously reported in skin *S. epidermidis* (22). While the majority of genes within the genus pan-genome were hypothetical, we detected multiple genes with a predicted role in successful skin colonization. Finally, we performed growth-and RNA-Seq analyses of select isolates from all three species in conditions that mimic the skin environment. Gene expression analysis revealed extensive variation between isolates, including within shared genes, highlighting the natural diversity in expression and possible regulatory mechanisms, which further contribute to the functional diversity in skin-associated staphylococci.

## RESULTS

### Healthy human skin harbors multiple *Staphylococcus* species

To identify staphylococcal species that are resident members of the human skin microbiome, we re-analyzed an extant skin metagenomic dataset obtained from fourteen body sites of twenty-two healthy volunteers that had previously been analyzed at the genus level (**Fig. S1** shows body sites sampled with abbreviations). The selected body sites encompassed multiple sebaceous, moist, and dry skin surfaces from head to toe. In agreement with previously published results, *Staphylococcus, Cutibacterium,* and *Corynebacterium* were the most prominent genera represented in the microbiome of these skin sites (23) (**Fig. S2**). The mean percent relative abundance of *Staphylococcus* varied widely across body sites from 4.3 ± 1.5% on back, to 64.1 ± 6.3% at the plantar heel. In general, *Staphylococcus* dominated the moist sites (37.3 ± 2.5%), where it was present at a significantly higher proportion than sebaceous sites (13.2 ± 1.4%; Wilcoxon rank-sum test, P value < 0.001) and dry sites (12.0 ± 2.1%; Wilcoxon rank-sum test, P value < 0.05). As this dataset interrogated the 5’ end of the 16S rRNA gene, there were sufficient high-confidence amplicon sequence variants (ASVs) exist to impute species level relative abundance with DADA2. Species that were present at ≥ 1% relative abundance at one or more body sites and were detected in more than three individuals were considered skin-resident species for this study (24). Using these criteria for abundance and prevalence thresholds, a total of seventeen staphylococcal species were identified as residents or indigenous to the human skin (**Fig. 1A**). Each body site was colonized simultaneously by multiple skin-resident staphylococcal species, with the average ranging from 2.6 ± 0.3 species to 5.3 ± 0.4 species on back and toenails, respectively.

**Figure 1.**
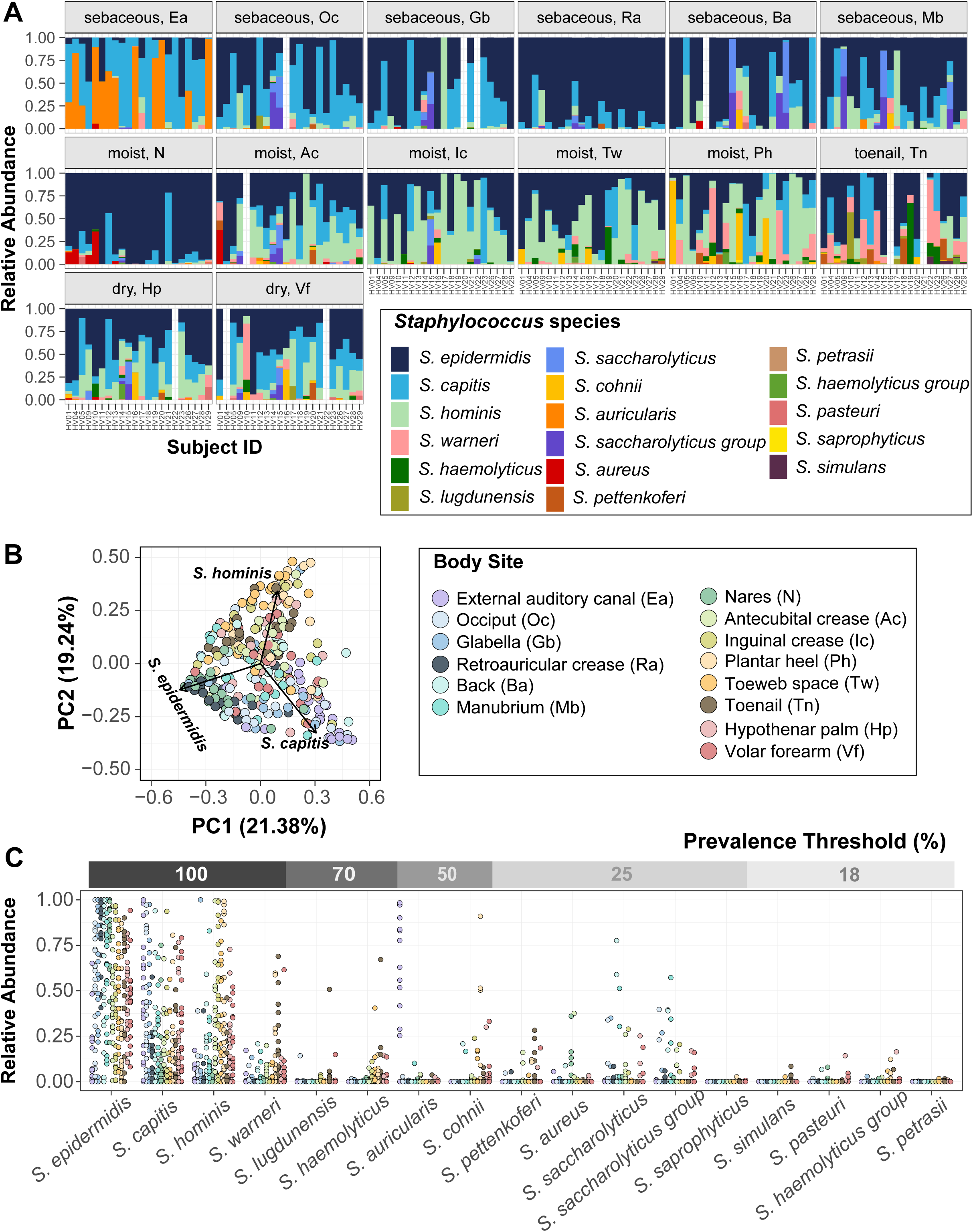
Diversity of staphylococcal species present on healthy human skin. **(A)** Barplots display the relative abundance of staphylococcal species at 14 body sites. Each color represents a distinct species as shown in the legend. Each bar represents one Subject/Healthy Volunteer (HV). Empty bars represent missing data. **(B)** Beta diversity analysis of staphylococcal communities present at different body sites using principal-coordinate analysis (PCoA) plot based on Bray-Curtis dissimilarity (PERMANOVA; R^2^ = 0.26804, P value < 0.0001). First two coordinates are shown accounting for 40.62% of the total variance. Individual staphylococcal ASVs driving the largest separation of body sites (displayed as colored dots) are shown as black arrows and are labeled by the corresponding species. Note that the body site legend is shared between figures B and C. **(C)** Relative abundance and prevalence of staphylococcal species detected in our dataset. Each dot represents proportion of a staphylococcal species relative to all other staphylococcal species in a sample. Dots are colored by body sites as shown in the shared legend. Percent prevalence threshold for a species to be considered part of the core staphylococcal community is shown in the upper gray bar (see text for details). Refer to Fig. S1 for body site details.

Some body sites displayed distinct staphylococcal community composition, characterized by a higher proportion of select species and indicating site preference. The most intimate relationship observed was the external auditory canal (Ea) where ∼55% healthy volunteers (12/22 Ea samples) were colonized by *S. auricularis* at a mean relative abundance (MRA) of 66.4 ± 7.8%. *S. auricularis* was rarely detected in other samples from these healthy volunteers (15/156 samples, 2.3 ± 0.6 MRA, Wilcoxon rank-sum test, P value < 0.0001 for MRA of *S. auricularis* in positive Ea samples versus all other positive samples). Further, barring HV14 (toenail sample with 0.04% MRA), *S. auricularis* was completely absent in individuals that lacked Ea colonization.

*S. epidermidis, S. capitis,* and *S. hominis* species were present in most samples (*S. epidermidis*: ∼ 95% samples at 52.3 ± 1.9 % MRA; *S. capitis*: ∼ 75% samples at 26.3 ± 1.8 % MRA; *S. hominis*: ∼ 70% samples at 23.7 ± 1.9 % MRA). However, even for these prominent species, colonization across body sites was not uniform. *S. epidermidis* predominated the nares (N) and retroauricular crease (Ra) (MRA: N: 83.7 ± 4.3 % and Ra: 86.0 ± 3.7 % as compared to all other sites: 46.3 ± 1.9 %; Wilcoxon rank-sum test, P value < 0.0001). *S. capitis* displayed a moderate site preference for the sebaceous head region (combined Ea/glabella/occiput: 45.4 ± 4.3 % versus non-head: 20.5 ± 1.7 %; Wilcoxon rank-sum test, P value < 0.0001), whereas *S. hominis* preferred moist sites (combined antecubital crease/inguinal crease/plantar heel/toeweb: 39.4 ± 3.4 % versus non-moist: 14.00 ± 1.6 %; Wilcoxon rank-sum test, P value < 0.0001). To assess if site preferences shaped staphylococcal community composition at different body sites, we calculated the Bray-Curtis dissimilarity. Unsupervised ordination of the dissimilarity matrix using principal coordinate analysis (PCoA) revealed separation of body sites largely driven by the three most prominent staphylococcal species, as indicated by labeled arrows on the plot (PERMANOVA; R^2^ = 0.27, P value < 0.0001; **Fig. 1B**). Specifically, the sebaceous head region and the moist sites were separated by *S. capitis* and *S. hominis*, respectively, whereas the nares and retroauricular crease were grouped together by *S. epidermidis* colonization.

We next sought to define a core staphylococcal community by prevalence threshold, based on proportion of healthy volunteers being colonized by a species, irrespective of relative abundance. (**Fig. 1C and Table S1)**. Of the seventeen skin-resident species, only *S. epidermidis, S. capitis, S. hominis, and S. warneri* were present on all volunteers, making them truly ubiquitous members of the skin microbiome. In addition to being most prevalent, *S. epidermidis, S. capitis*, and *S. hominis* also showed the highest MRA and could be detected at all skin sites in highly variable proportions (**Fig. S3**). Decreasing the prevalence threshold below 100%, increased the number of species that can be considered core, with all species being included at a threshold of 18%; however, most of these low prevalence species, including *S. aureus*, were also present at low relative abundance and detected only at select body sites (**Fig. 1C**).

Collectively, our skin metagenomic sequencing analyses revealed diverse staphylococcal communities with body site specific compositions. Given the predominance of *S. epidermidis*, *S. capitis,* and *S. hominis* on the skin, we focused on these three species for in-depth genomic and functional analyses.

### Genome sequencing captures intra-species diversity

We sought to catalog the genetic makeup and diversity of human skin-associated staphylococci by concentrating on *S. epidermidis, S. capitis*, *and S. hominis* genomes by leveraging our extensive skin microbiome culture collection, which contains diverse microbial isolates from multiple body sites of healthy volunteers. A total of 273 staphylococcal isolates were sequenced using Illumina short reads, after an initial taxonomic classification by species-specific PCR and full-length 16S rRNA gene sequencing. After removing highly similar genomes by de-replication (> 99.9% ANI), our final list consisted of 22 *S. capitis* (9 volunteers, 10 body sites), 49 *S. epidermidis* (12 volunteers, 15 body sites) and 55 *S. hominis* (11 volunteers, 12 body sites) non-clonal complete genomes (N = 126 total) (**<u>Fig. S1, Table S2</u>**). fastANI values of de-replicated genomes revealed sharp boundaries separating species as shown in **<u>Fig. S4</u>** (25). Comparison of key genome statistics showed that *S. hominis* isolates had the smallest average genome size and the least number of predicted protein coding genes (**Table 1, <u>Table S2)</u>** We confirmed that the *S. hominis* result is not restricted to our dataset by analyzing publicly available genomes (**Fig. S5**), which also demonstrated a genome size smaller than other sequenced staphylococcal species.

**Table 1.** Genome characteristics of each species based on isolates used in this study.

| **Genome characteristic** | ***S. epidermidis***<br>**(N = 49)** | ***S. capitis***<br>**(N = 22)** | ***S. hominis***<br>**(N = 55)** |
| --- | --- | --- | --- |
| Mean Genome Size (Mb) | 2.507317 ±<br>0.085426 | 2.480510 ±<br>0.049336 | 2.273876 ±<br>0.077851* |
| Mean GC-content (%) | 36.7 ± 0.2 | 36.9 ± 0.3 | 36.2 ± 0.2 |
| Mean number of<br>protein coding genes (CDS) | 2315 ± 88 | 2378 ± 47 | 2195 ± 93** |

\*, \*\* *S. hominis* has a significantly smaller genome and encodes fewer CDSs than the other two species surveyed (p < 0.01, one-way ANOVA, post hoc Tukey's HSD)

### Species-and genus-level pan-genome analysis reveals extensive genetic diversity

Using a pan-genomics approach, we identified orthologs and clustered genes conserved across all three species versus those that were taxonomically restricted to one species or even a subset of strains. To compare the protein coding gene content across species, we initially calculated three independent species-level pan-genomes to generate a non-redundant catalog of all protein-coding genes (CDS), along with their distribution within conspecific isolates (**Table 2, Table S3**).

**Table 2.**
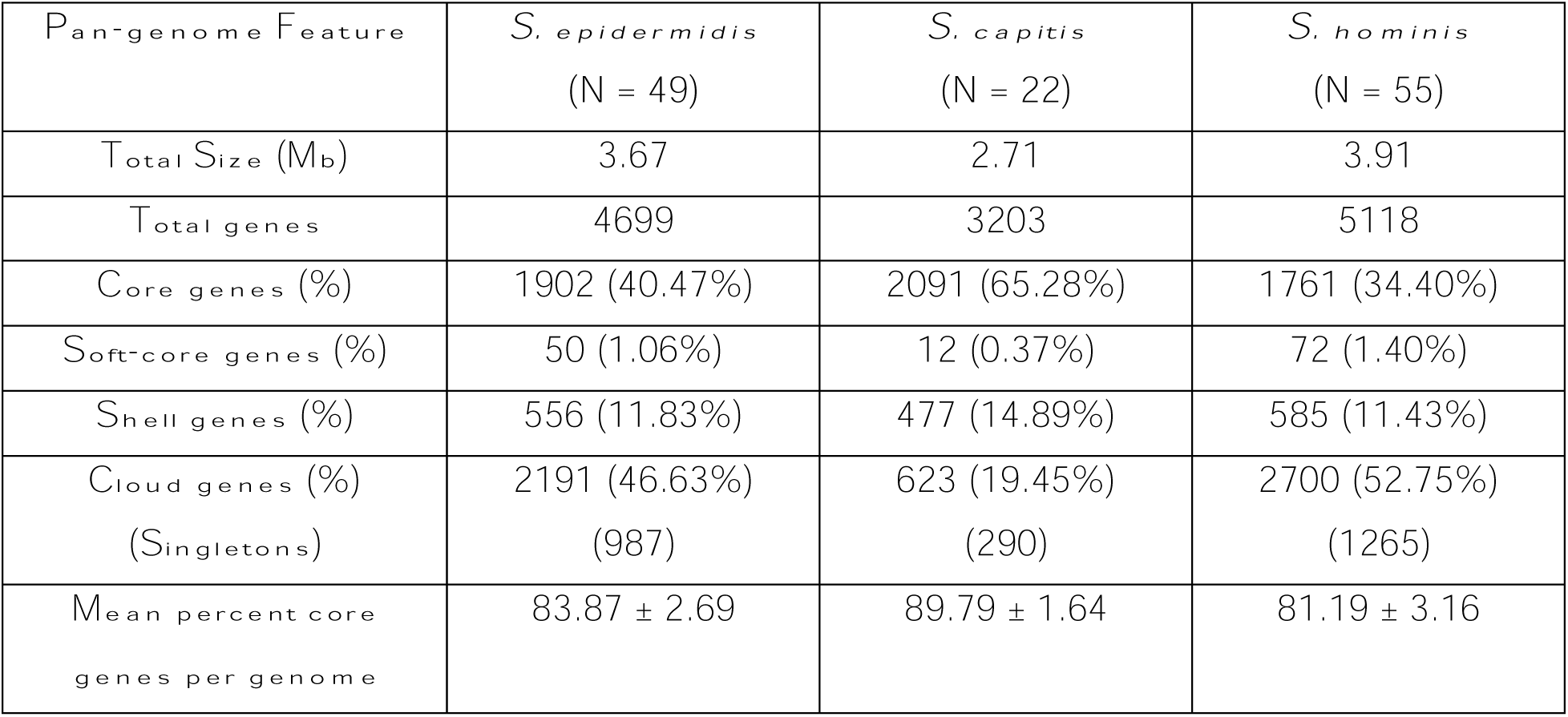
Major pan-genome features of each species.

| **Pan-genome Feature** | ***S. epidermidis***<br>**(N = 49)** | ***S. capitis***<br>**(N = 22)** | ***S. hominis***<br>**(N = 55)** |
| --- | --- | --- | --- |
| Total Size (Mb) | 3.67 | 2.71 | 3.91 |
| Total genes | 4699 | 3203 | 5118 |
| Core genes (%) | 1902 (40.47%) | 2091 (65.28%) | 1761 (34.40%) |
| Soft-core genes (%) | 50 (1.06%) | 12 (0.37%) | 72 (1.40%) |
| Shell genes (%) | 556 (11.83%) | 477 (14.89%) | 585 (11.43%) |
| Cloud genes (%)<br>(Singletons) | 2191 (46.63%)<br>(987) | 623 (19.45%)<br>(290) | 2700 (52.75%)<br>(1265) |
| Mean percent core<br>genes per genome | 83.87 ± 2.69 | 89.79 ± 1.64 | 81.19 ± 3.16 |

We then used a reciprocal-best-blast-hits approach to cluster orthologs from species pan-genomes and created a merged genus-level pan-genome with core and accessory gene partitions (**Table S4**). The pan-genome matrix consisted of 7744 representative genes shared across all staphylococcal genomes, of which 21% (1570 genus-core + 90 genus-soft-core genes; total 1660 genes) were found in ≥ 95% genomes and were termed genus-core, while the remaining 79% (1116 genus-shell genes + 4968 genus-cloud genes, including 1717 singletons) were accessory genes present anywhere between 1 to < 95% of all genomes (**Fig. 2A**). Most accessory genes were restricted to 1-2 species (**Fig. 2B**). Interestingly, even within the accessory partition, a sizable proportion of genes (∼11%; 683/6084) were core to individual species (present in ≥ 95% of species genomes), indicating species-specific selection pressure to retain them (**Fig. 2B**). Distribution of genes within each genome, along with their sharing between genomes was demonstrated by a gene presence-absence matrix (**Fig. 2C)**; sharing was quantified by Jaccard index measuring pair-wise gene sharing between all genomes (**Fig.2D**). Since the genus-core of 1660 genes accounted for ∼70% of protein coding genes per genome (total: ∼2300 CDS/genome based on Table 2), this indicated that most of the genes encoded within a genome are conserved between all species.

**Figure 2.**
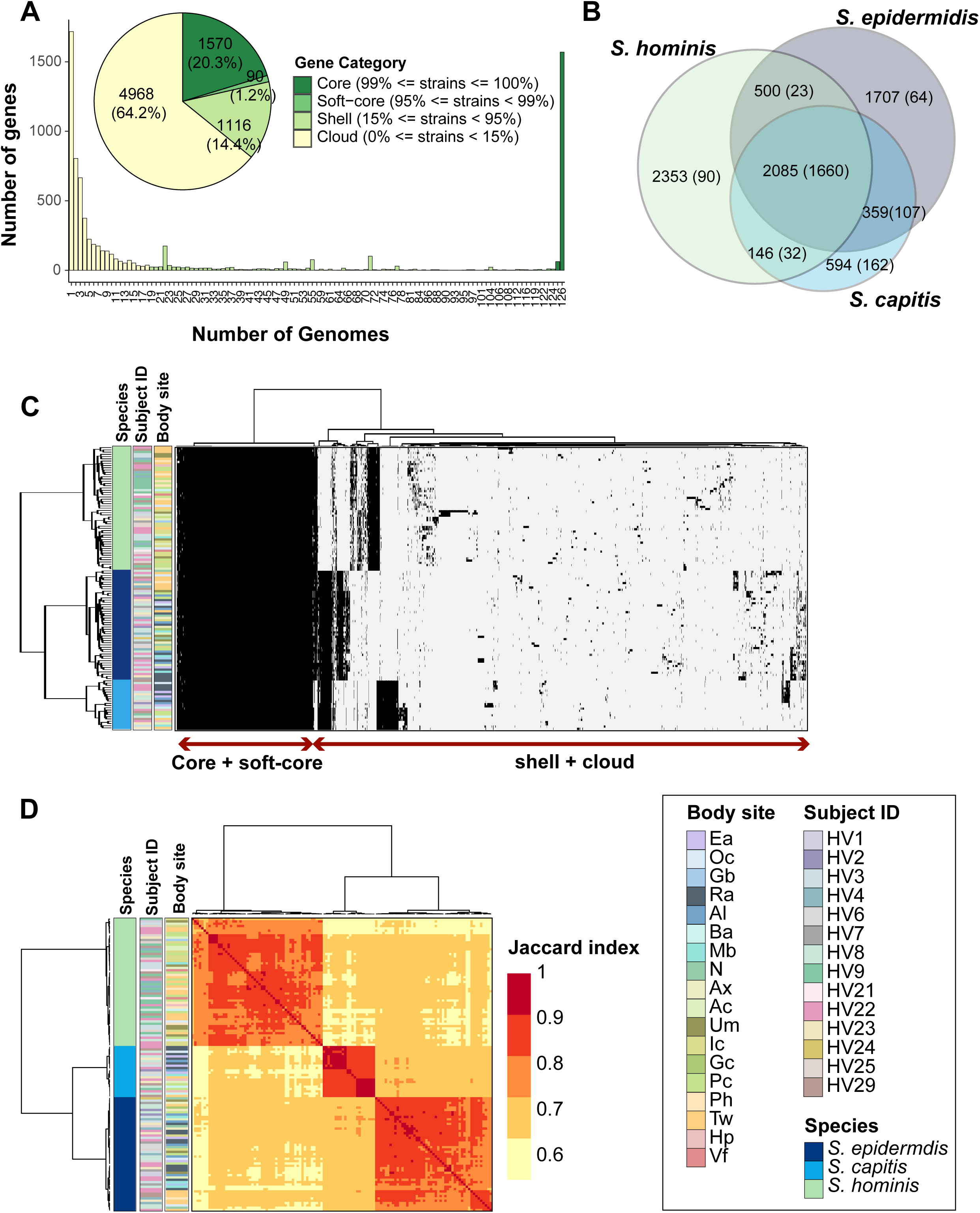
Genus-level pan-genome summary to quantify gene sharing between species. **(A)** Histogram of core, soft-core, shell, and cloud genes of the genus-pangenome. Pie chart shows the number of genes in each category as defined in the legend; percent of genes in each category are displayed in brackets. **(B)** Venn diagram displaying the distribution of 7744 genes between the three species. Number of total genes in each intersection are shown outside the bracket and core genes in each intersection (genes detected in ≥ 95% genomes present in the intersection set) are shown inside brackets. **(C)** Clustering of all 126 genomes using Euclidean distance and Ward hierarchical clustering based on the presence/absence patterns of 7744 genes detected in the pan-genome. Sidebars display species, volunteer, and body site of isolation for each genome. **(D)** Pairwise Jaccard index between genomes displaying the proportion of shared gene content. Note: Distance of 1.0 depicts complete gene overlap, with lower numbers representing lesser degree of gene sharing. Hierarchical clustering of genomes and the displayed sidebars are the same as that shown in figure C. Note that the legend is shared between figures C and D. Refer to Fig. S1 for body site details.

### Functional analysis shows conserved core metabolic functions between species

We next sought to annotate the genus pan-genome to dissect functional capabilities that are central to all species, versus those that are species restricted. The percentage of genes that had an annotation or a predicted Pfam domain was ∼63%, and this percentage progressively decreased from core (93%) to shell (79%) to cloud (49%) genes (**<u>Fig. S6A</u>**). A comparison of the top 20 most represented Pfam domains in each pan-genomic category revealed considerable differences between core, shell, and cloud fractions (**<u>Fig. S6B</u>**). Specifically, the core gene’s domains were involved in transport and central metabolic enzymes, while the annotated cloud genes predominantly carried domains found in phages, transposable elements, and restriction modification systems. None of the Pfam domains were exclusive to or enriched in any one species (data not shown). Over-representation of canonical metabolic pathways within the genus-core was also supported by KEGG enrichment analysis (**<u>Fig. S6C</u>**). This suggested that all species have similar central metabolic capabilities, which was confirmed by mapping genes onto KEGG modules to determine completeness of canonical pathways (**Fig. 3**). Using a strict criterion of ≥ 75% module completeness in at least one genome, we found sixty-two modules, of which forty-six were complete in all the genomes, irrespective of species. Three complete core metabolic modules were present in individual species and were not part of the genus-core. These included M00026 (histidine biosynthesis) in *S. epidermidis*, M00045 (histidine degradation) in *S. capitis*, and M00127 (thiamine biosynthesis) in *S. hominis* genomes.

**Figure 3.**
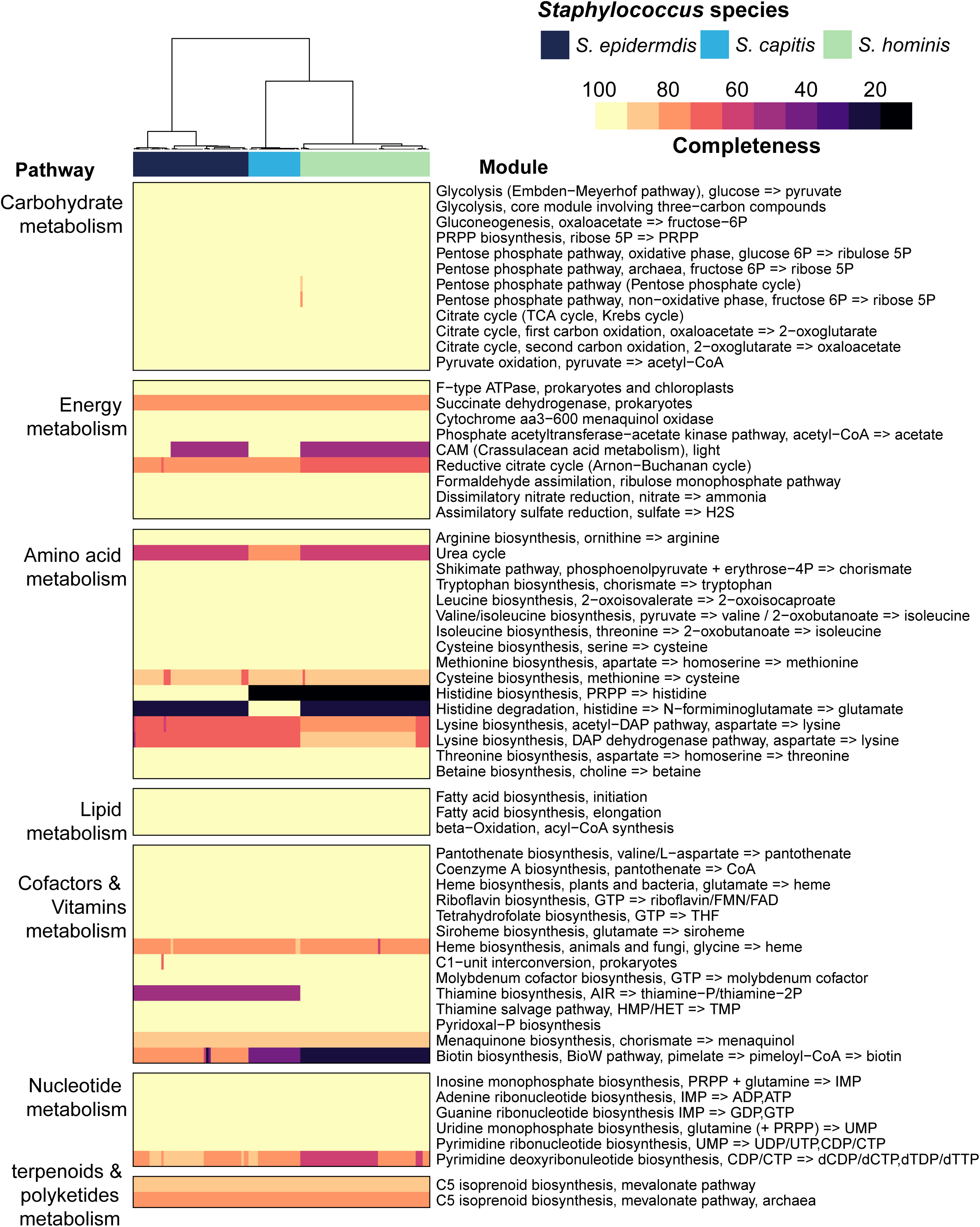
Core metabolic pathways present in staphylococcal genomes using KEGG module analysis. Completeness of a pathway is expressed as a percentage from 0 to 100. Only pathways that showed ³ 75% module completeness in at least one genome are shown. Hierarchical clustering (Euclidean distance; Ward) of species is shown as a dendrogram. Details of modules can be found at: https://www.genome.jp/brite/ko00002

Along with these forty-six metabolic pathways, the genus-core also carried several genes/operons with a predicted function in skin colonization including immune evasion (*srtA, dltABCD, mprF, sepA, vraFG, vraX, graSRX, capABC, oatA)*, acid response (*ureABCEFGD, yut*), osmotic tolerance (*betA, opuCABCD*), biofilm formation (*atlE, pmtABCD*), iron acquisition (*yfmCDE, isdG*), and niche occupancy (*tagDXBGHA, gehC*) (**Table S5**).

### Species-restricted core-genes carry loci with predicted role in skin colonization

We next analyzed the 683 taxonomically restricted core genes within the accessory partition that were present as core in 1-2 species (≥ 95% species genomes), and they served as candidates under strong selection for conferring unique capabilities to individual species (**Table S6**). These included 64, 162, and 90 genes that were unique to *S. epidermidis*, *S. capitis,* and *S. hominis* respectively, 107, 32, and 23 genes that were shared by all isolates of *S. epidermidis* & *S. capitis*, *S. epidermidis* & *S. hominis*, and *S. capitis & S. hominis*, respectively, and 205 genes that in addition to being core to 1-2 species, were also detected at a lower frequency in remainder genomes (**Fig. 2B**). We focused on annotated genes, especially ones that play a role within the skin environment (either by themselves or in combination with genus-core genes, **Fig. 4, Table S6)**. Some genes were present as paralogs of genus-core genes, with potentially divergent functionality. For example, the D-and L-methionine transport locus (*metNIQ*) was present as part of the genus-core, with an additional copy in all *S. epidermidis* genomes. Interestingly, some of the species-restricted-core genes were also present at a lower frequency in other species, indicating species-level differences in selection pressure to retain these genes. Of note is the biosynthetic pathway for the broad-spectrum metallophore, staphylopine, that was present in all *S. epidermidis* and many *S. capitis* (N = 16/22) genomes (26). The pathway requires *L-histidine*, which can either be synthesized by all *S. epidermidis* isolates (KEGG module M00026) or can be imported by *S. capitis* species (KEGG module M00045). Ni^2+^ is imported with high affinity by staphylopine, which then acts as a urease co-factor (genus-core genes) and plays an important role in survival within the acidic skin environment (27). Another well-known example is the *icaABCDR* locus involved in biofilm formation, that was core to *S. capitis*, and accessory to *S. epidermidis* (N = 17/49), where it is considered to be a virulence factor (28). Besides these examples, fitness determinants involved in resistance to reactive oxidants, host generated polyamines, and antimicrobial fatty acids, acid tolerance, biofilm formation, resistance to antibiotics, and osmoregulation were present as species-restricted-core genes as shown in Fig. 4. Metabolic genes involved in fermentation pathways (glycerol, lactate, formate (29), and acetoin metabolism) that are important under glucose limiting and microaerobic skin conditions, and secreted proteases (metalloproteases: SepA (30), cysteine protease Ecp and serine protease Esp (31), and a sphingomyelinase (*sph* (11)) that play a role in microbe-microbe and host-microbe interactions were also detected. Well-characterized genes that were restricted to either *S. capitis* or *S. hominis* were rare, which suggests novel functions encoded by these species. One such example is the *patB* gene present in all *S. hominis* genomes that was recently shown to be responsible for body odor-associated thioalcohol generation from the odorless precursor Cys-Gly-3M3SH secreted by apocrine glands (32). Although the biological role of thioalcohol generation is currently unknown, the inguinal crease is known to harbor a high density of apocrine glands, a preferred site for *S. hominis* colonization.

**Figure 4.**
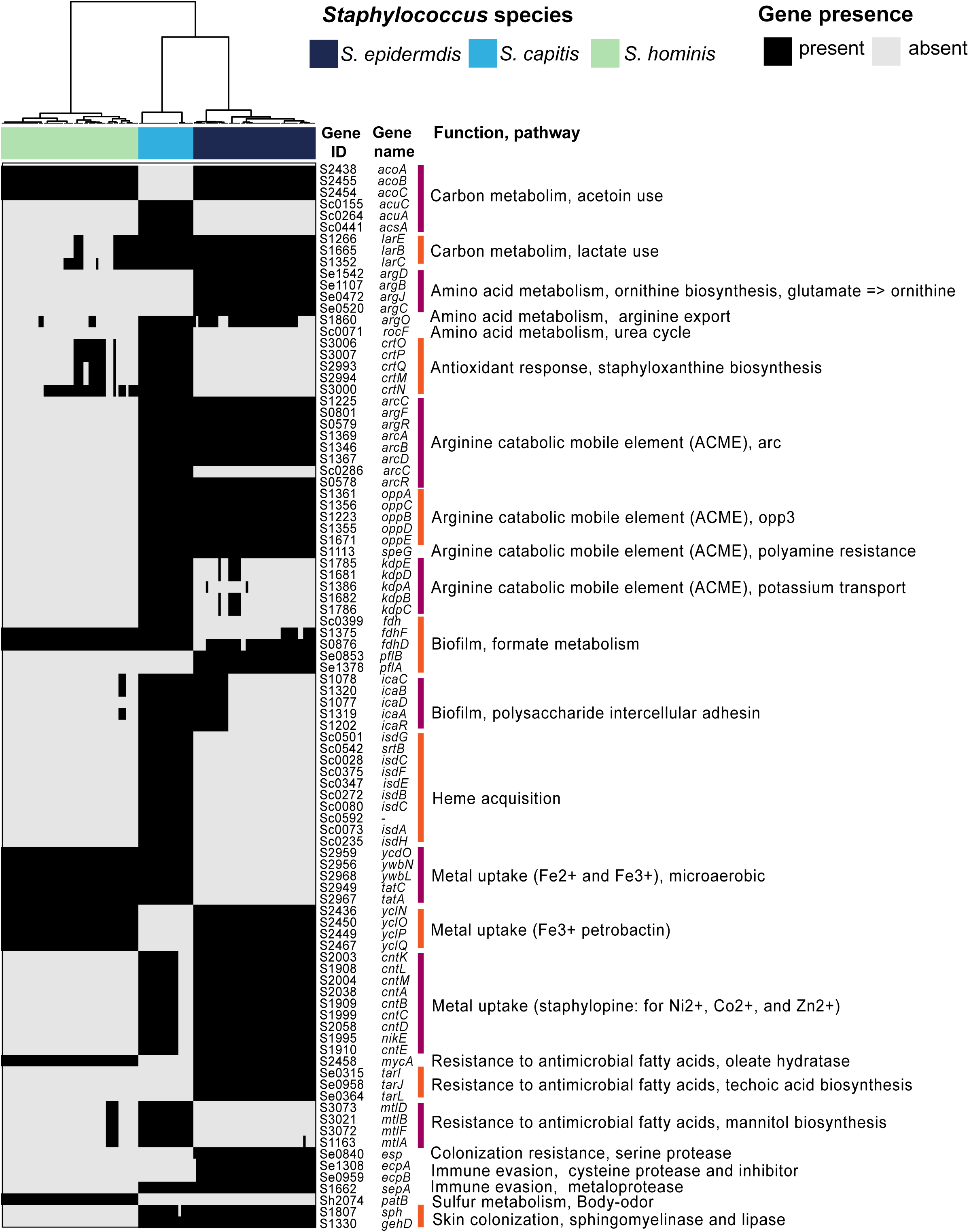
Species-restricted core genes relevant to skin colonization. Presence/absence patterns of select genes/loci that encode a complete pathway (by themselves or along with genus-core genes) and have a known role in skin colonization are shown. Gene ID, gene names and function encoded by each gene is shown on the right. Hierarchical clustering (Euclidean distance; Ward) of species is shown as a dendrogram on top.

Apart from the species-restricted-core genes, majority of the accessory genes (N = 5401; ∼68%) were variably distributed in a subset of isolates or were singletons, likely conferring strain-specific functions. Most of these genes were uncharacterized with no functional annotation or encoded phage-and transposon-like functions.

### Intra-species phylogenetic clades display distinct gene sharing and body site preferences

Studies in other microbes have shown that the degree of gene sharing between two strains of the same species is primarily dictated by their evolutionary relatedness (33). To test this hypothesis in *Staphylococcus*, we carried out phylogenetic analysis of conspecific genomes using a maximum-likelihood tree based on nucleotide polymorphisms in core genes alignments. A genus-level phylogenetic tree based on an alignment of select genus-core genes (N = 665) was generated to root individual species trees (**Fig. S7)**. In all three species, phylogenetic analysis revealed the presence of two divergent phylogenetic clades with 100% bootstrap support (**Fig. 5A-C**). Grouping of isolates based on the pan-genome was tested by hierarchical clustering using the Jaccard distance (**Fig. 5A-C**), and by principal component analysis (PCA) (**Fig. S8**). This identified groups that broadly corresponded with the phylogenetic clades, and the overall pairwise pan-genomic distance within a clade was lower than between clades (**Fig. 5D**).

**Figure 5.**
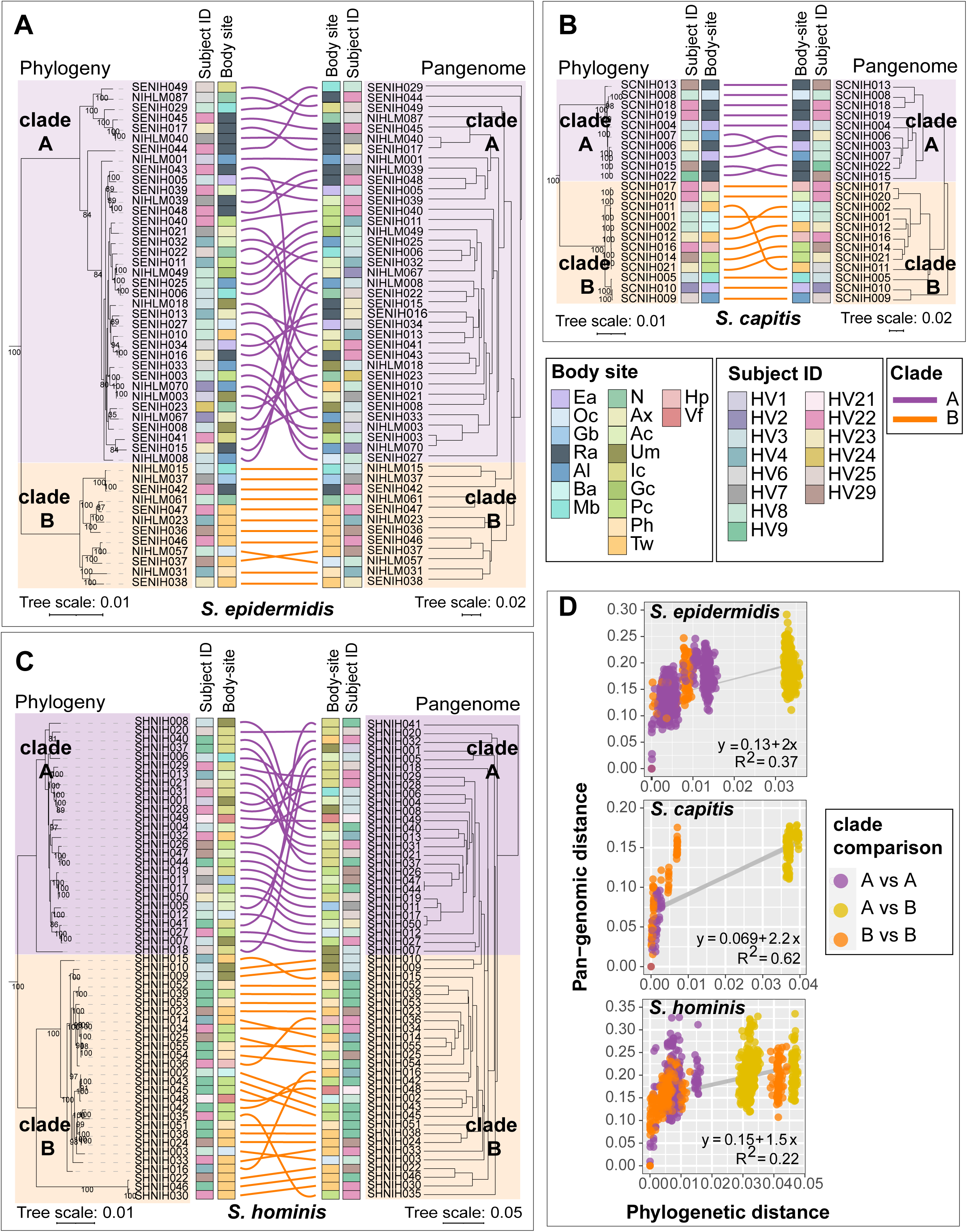
Relationship between phylogeny and pan-genome in conspecific staphylococcal genomes. **(A-C)** Tanglegrams depicting the relationship between the phylogenetic tree based on core genes sequence alignment (left) and hierarchical clustering (binary distance; average linkage) of the pan-genome presence/absence matrix (right). Clades A and B for each species are shown in purple and orange color, respectively. Colored lines are used to connect the same isolate on two trees of the tanglegram. Healthy volunteer and body site of isolation for each isolate are displayed as colored strips within the tanglegram. **(D)** Correlation between phylogenetic distance and pan-genome distance for all pair-wise genome comparisons within a species. Each dot represents one pair-wise comparison. Dot colors represent the clades to which the genomes being compared belong, with yellow color dots representing clade A vs clade B comparison. Refer to Fig. S1 for body site details.

A pan-GWAS analysis was used identify clade-specific gene enrichment in all three species (**Table S7**). For *S. epidermidis* and *S. capitis* the results were concordant with previously published pan-genomes (17, 28, 34). Namely, formate dehydrogenase gene (*fdhH*) involved in formaldehyde detoxification was exclusively present in all *S. epidermidis* clade B isolates, whereas clade A, which covered isolates from bloodstream infections in earlier studies, was enriched for genes involved in host cell adhesion, antibiotic resistance, biofilm formation, and virulence genes. For *S. capitis*, the clades corresponded with *S. capitis* subspecies, specifically, the commensal *S. capitis* subspecies *capitis* (overlap with clade A), and the more pathogenic *S. capitis* subspecies *ureolyticus* (overlap with clade B). All clade A isolates carried the arginine catabolic mobile element (ACME), a histidine decarboxylase (*hdcA*) that synthesizes the neurotransmitter histamine, and Type I restriction-modification system (*hsdRM*). Clade B isolates were enriched for genes involved in the staphylopine biosynthesis (*cntKLM*), a L-malate dehydrogenase (*mqo*), and *hxlAB* genes encoding formaldehyde detoxification system. For *S. hominis*, clade A encoded *dhaKLM* genes important for anaerobic utilization of glycerol, which is abundant on human skin, while clade B showed enrichment of lactate racemase (*larABCDE*), allowing the use of L-and D-lactate that is found in abundance at moist skin sites.

Intriguingly, clades showed significant enrichment by body sites (Fisher’s exact test, P value < 0.05), indicating site or habitat preferences. *S. epidermidis* clade B was enriched for feet sites (7/12) compared to clade A (1/36), which did not show any habitat preference (**Fig. 5A**) In *S. capitis* all clade A isolates were cultured from head sites (10/10), a region known to be the primary habitat of *S. capitis* colonization (35). On the other hand, *S. capitis* clade B isolates were cultured from diverse skin sites, with only 2/12 coming from the head region (**Fig. 5B**). For *S. hominis* each clade was enriched for a different body site, with clade A showing an over-representation of inguinal crease isolates (10/27 in A vs 0/28 in B), and clade B of feet region isolates (1/27 in A vs 15/28 in B).

Overall, our comparative pan-genome analysis between all three species revealed impressive degree of gene conservation across species. Further, we detected many skin-relevant functions in the taxonomically restricted core genes, indicating genetic traits for successful skin colonization that were unique to each species. In addition to these well-conserved genes, the accessory genes, with several known fitness determinants, most likely represent clade-driven niche-specialization that allow colonization of specific body sites.

### Growth curves in skin-like media conditions reveal intra-species phenotypic diversity

In addition to pan-genomic differences, we were interested in identifying gene expression differences that may exist between staphylococcal isolates under similar skin-like growth conditions. We specifically wanted to test if the shared genes showed similar expression patterns, with accessory and unique genes driving the major gene expression differences between isolates.

We first compared the ability of our isolates to grow in conditions that mimic the skin environment, which can be broadly defined as acidic and nutrient-poor, with amino acids and lipids as the main energy sources (36). We tested an isolate collection (n = 11 to 16) from each species, which represented ³ 88% of the species pan-genome (without singletons). Kinetic growth analysis was carried out in two artificial skin media representing the acidic, low nutrient (Eccrine sweat; ES), and the sebaceous or oily (Eccrine sweat with lipids; ESL) conditions present on healthy human skin (**Table S8**). Brain heart infusion supplemented with yeast extract (BHI-YE) was used as a control medium, which supported robust growth of all isolates (**Fig. 6A-C**). At the end of 24 hours, we observed considerable intra-species growth variation in nutrient-limited ES and ESL media, as quantified by area under the curve (AUC) (**Fig. 6D-F)**. Addition of lipids slightly improved the mean AUC for all species (1.3-, 1.5-and 1.1-fold for *S. epidermidis*, *S. capitis*, and *S. hominis*, respectively), but was not statistically significant. We did not observe any clade-specific growth differences in *S. epidermidis* and *S. hominis* isolates. In contrast, clade B isolates of *S. capitis* grew consistently better than clade A isolates in all media (P value < 0.001; t-tests comparing mean AUC for clade A vs clade B isolates, independently in each medium), suggesting divergent growth requirements between the two phylogenetic clades.

**Figure 6:**
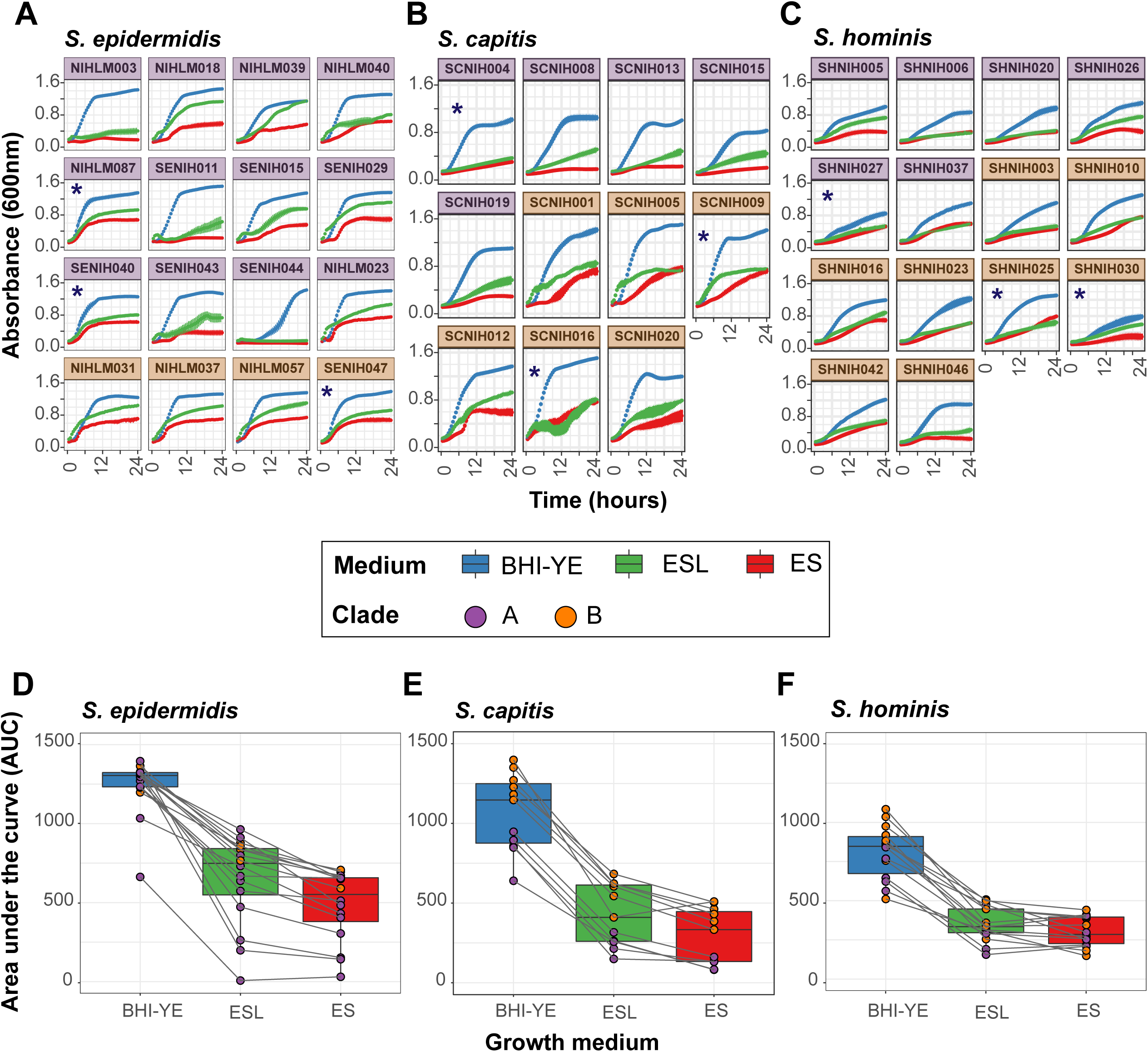
Kinetic growth analyses of select staphylococcal isolates in BHI-YE and artificial skin media, ES and ESL. **(A-C)** Growth curves of select *S. epidermidis*, *S. capitis*, and *S. hominis* isolates grown independently in three culture media, as measured by absorbance at 600nm over 24 hours. Each growth curve is an average of four independent biological replicates; error bars for each curve are shown. Isolates marked with an asterisk (*) were selected for RNA-Seq analyses. Purple and orange color of facets indicates the phylogenetic clade to which each isolate belongs. Growth curve color represents the growth medium. **(D-F)** The area under the curve (AUC) was used to quantify the growth of each isolate shown in A-C. Boxplots depict combined AUC values for all isolates of a species in each growth medium, as depicted by the color of the boxplot. The center black line within each boxplot represents the median value, with edges showing the first and third quartiles. Individual AUC values are shown as dots colored by clade of the isolate. Note that the legend is shared between all figures.

### RNA-Sequencing analysis shows considerable variation in gene expression between all isolates

Based on our growth curve analysis, we chose three isolates from each species for RNA-sequencing (RNA-Seq) (shown with an asterisk in **Fig. 6A-C**), such that they belonged to different clades/sub-clades, and whenever possible, showed distinct growth phenotypes in our skin media. Each isolate was grown separately in triplicate in BHI-YE, ES, and ESL medium, and cells were harvested at mid-log phase for RNA-Seq.

To make preliminary comparisons between isolates in all the three media, we restricted our initial RNA-Seq analysis to 1647 genus-core genes that had at least one read in all samples. This sub-setting was done to prevent any clustering bias due to differences in gene presence/absence. A principal component analysis (PCA) (**Fig. 7A**) and hierarchal clustering (**Fig. S9**) of variance stabilized read counts from the resulting dataset showed distinct clustering of BHI-YE samples from skin-like media. Further, samples from each medium grouped together by species, with *S. hominis* samples being more distant than the other two species. Given the overlapping profiles of ES and ESL samples for each species, only the ES samples were further analyzed.

**Figure 7:**
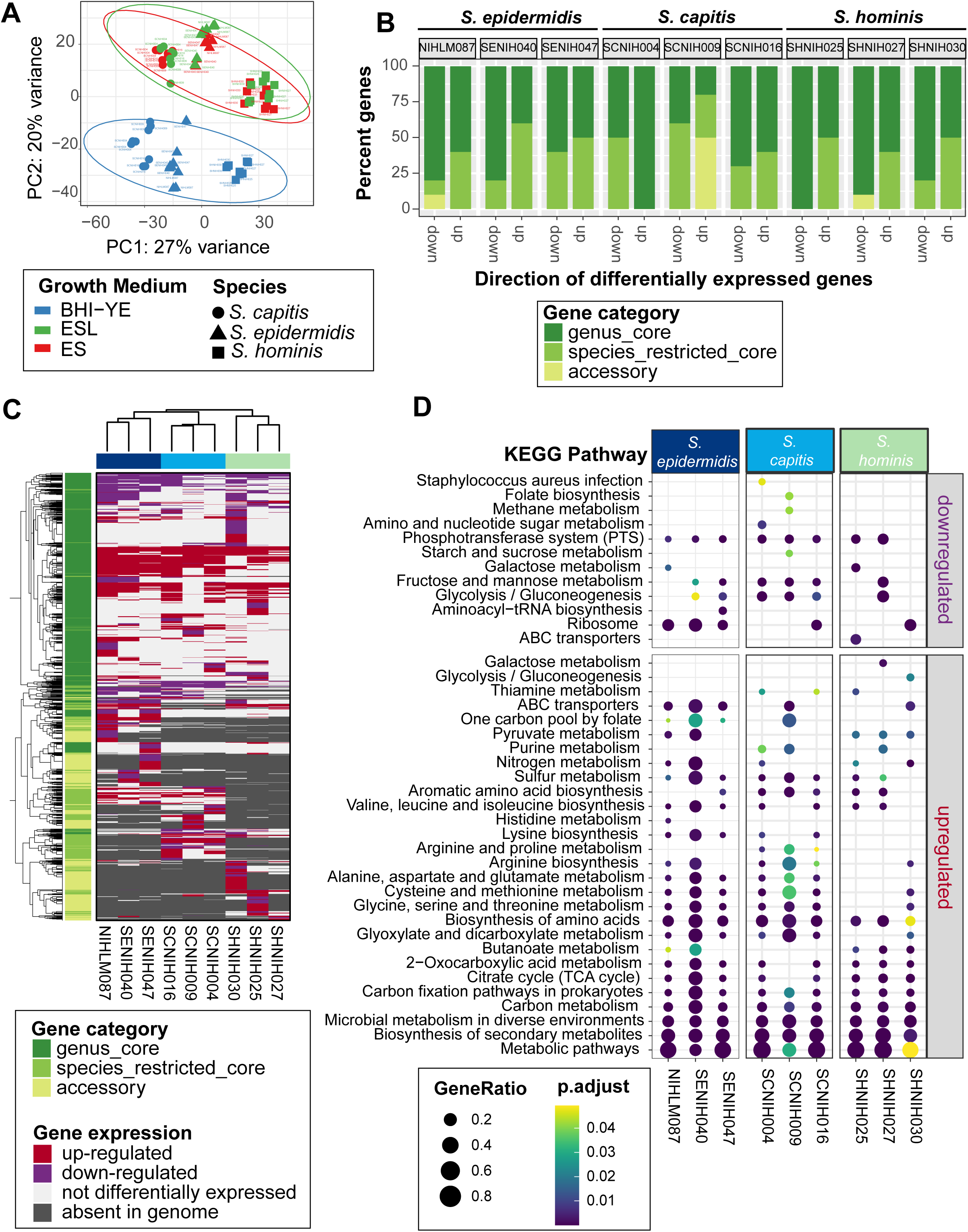
RNA-Seq analysis of nine distinct staphylococcal isolates in BHI-YE and in artificial skin media, ES and ESL (N = 3 biological replicates). **(A)** Principal-component analysis of variance stabilizing transformation-normalized reads from all RNA-seq samples and 1647 genus-core genes. Colors represent different growth media, and shapes represents staphylococcal species. **(B)** A stacked barplot representing the proportion of genus-core, species-core, and accessory within the top 10 down-and up-regulated genes in each isolate in the ES medium relative to BHI-YE. **(C)** A discrete heatmap of log_2_-fold changes of all the genes that were differentially expressed (≥ or ≤ 2-fold change, adjusted P value < 0.05) in ES relative to BHI-YE. Columns are individual isolates and rows are differentially expressed genes. Genes upregulated in the ES medium are shown in red, downregulated genes are in purple. Light pink cells represent genes that were encoded within an isolate genome, but not differentially expressed in that isolate. Grey cells denote genes that were absent in the isolate genome. The left sidebar depicts the category to which each gene belongs. Top and side dendrograms were generated by unsupervised clustering of the expression data (Euclidean distance; Ward). **(D)** KEGG pathway enrichment analysis of differentially expressed genes in ES medium relative to BHI-YE in each isolate. The x-axis and y-axis show the isolate and pathway, respectively. Bubble size shows GeneRatio and color indicates adjusted P value.

To measure the contribution of species-specific core and accessory genes, in addition to the genus-core genes, we next quantified the differential gene expression of each isolate individually. The number of differentially expressed genes (DEGs, ≥ or ≤ 2-fold change; adjusted P value < 0.05) in ES medium relative to BHI-YE, and their distribution between the genus-core, species-restricted core and accessory gene categories for each isolate is shown in **Fig. S10 and Table S9-S10.** The majority of DEGs for each isolate belonged to the genus-core (median: 70%, range: 58%-74%), followed by species-restricted core (median: 18%, range: 13%-29%) and then accessory (median: 13%, range: 3%-17%), indicating that bulk of the gene expression differences between isolates lies within the conserved gene content. However, a closer look at the top ten upregulated and downregulated genes in the ES medium relative to BHI-YE in each isolate revealed that a disproportionately high percentage of upregulated genes (median 50%) belonged to the species-restricted core or accessory genes (**Fig. 7B**). Further, a comparison of gene expression patterns between all the nine isolates in the ES medium relative to BHI-YE revealed extensive variation in all gene partitions. A discrete heatmap displaying all 1979 DEGs (∼26% genus pan-genome) across all isolates is shown in **Fig. 7C**. We identified 145, 216, and 102 genus-core genes and 58, 50, and 18 species-restricted genes that were differentially expressed in the ES medium relative to BHI-YE by all three isolates of *S. epidermidis, S. capitis*, and *S. hominis*, respectively. This included upregulation of core central metabolism genes involved in gluconeogenesis (*pckA, gapB, ppdK, aldA, S0553*), tricarboxylic acid (TCA) cycle (*sdhA, sdhC, citB, citZ, acnA*), amino acid (*metB, metC, yitJ, rocA, rocD, putA, ilvB, yfmJ*) and nucleotide (*purE*) metabolism, competence transcription factor (*comK*), and an alpha-glucosidase (*yugT*) in all isolates. This was accompanied by downregulation of core genes involved in glycolysis (*cggR, pgk, gapA, fruA, fruK, ptsG*) and zinc transport (*adcA*) in the ES medium. Within the species-restricted core genes all *S. capitis* isolates showed upregulation of the *lar* operon, which is involved in utilization of L-and D-lactate that was present as the sole carbon source in the ES medium. Genes responding to the acidic conditions (pH 5.5) in the ES medium were upregulated in all *S. epidermidis* isolates, including *oppABCDF,* ornithine biosynthesis: *arcA, arcB, argJBCD*. In addition, genes involved in acetoin metabolism (*lpdA* and *acoABC*) that were core to *S. epidermidis* and *S. hominis,* but not *S. capitis*, were the most highly upregulated genes in all six isolates (median 254-fold change, range ∼30-to 2500), showing the importance of this pathway within the skin environment. Beyond these examples, however, there was a wide variation in expression of genes irrespective of the gene category. For example, mannitol metabolism pathway (*mltABDF*; > 4 fold change), a fitness determinant under hyperosmotic conditions present on the skin (37), was encoded by all *S. capitis*, but only expressed by SCNIH004 in the ES medium. On the other hand, *kdp* operon (*kdpABC*; 3-8 fold change), another *S. capitis* core locus that is induced by high NaCl concentration (38), was upregulated in the ES medium SCNIH009 and SCNIH016, but not in SCNIH004. Finally, we sought to examine the broad functional classifications of genes that are significantly enriched in the ES medium relative to BHI-YE. KEGG enrichment analysis of DEGs identified overall enrichment in global metabolism pathways, including carbon and amino acid metabolism in all isolates (**Fig. 7D**). Despite the variation in DEGs, we saw a consistent enrichment in several amino acid metabolism pathways in *S. epidermidis* and *S. capitis* isolates, and butanoate metabolism, which encompasses acetoin metabolism, was enriched in all *S. hominis* and two *S. epidermidis* isolates.

Collectively, our data revealed a species-level signature to gene expression, accompanied with an extensive variability between isolates of the same species, which could be largely attributed to differential expression in both, the genus-and species-core genes.

## DISCUSSION

The present study combines metagenomic, pan-genomic, and gene expression analyses for a detailed taxonomic and functional characterization of the most prevalent skin-associated staphylococci. By using the 16S rRNA gene as a taxonomic marker for species classification, our work revealed predominance of three staphylococcal species, *viz*. *S. epidermidis*, *S. capitis* and *S. hominis*, for which humans serve as the primary host (21). In addition, many other clinically relevant species, including *S. aureus*, *S. lugdunensis, S. pettenkoferi*, *S. haemolyticus, S. cohnii* and *S. saprophyticus* were also present in multiple volunteers at a lower relative abundance. Further, *S. aureus* reached its highest relative abundance in the nares, where it was detected in 22% individuals, in agreement with known frequency of *S. aureus* nasal carriage (39).

Our amplicon data suggested that *S. epidermidis*, *S. capitis,* and *S. hominis* were generalists, colonizing multiple body sites at variable frequencies. However, a pan-genome analysis using isolates with detailed provenance tracking (both volunteer and body site of isolation) revealed that each species contained isolates belonging to divergent phylogenetic clades with marked body site preferences, often co-colonizing the same healthy volunteer. Each clade was associated with distinct accessory genes, many of them encoding skin-relevant functions.

Stable co-existence of genotypic clusters from the same species is thought to arise by adaptation to novel resources with a non-uniform distribution within an environment (40). In-line with this hypothesis, human skin shows variations in temperature, pH, compositional differences in lipids, and in the expression of keratin and immune markers across different body sites (41), strongly suggesting the presence of spatially partitioned resources and anti-microbial factors, both determining strain colonization. Future work directed towards identifying genes that are under selection in distinct staphylococcal clades should shed light on molecular mechanisms shaping body site preferences in these isolates.

Comparative pan-genome analysis between all three species showed that, except for the essential and central metabolic genes, most of the genes (79%) were not uniformly shared across all species. One caveat of our pan-genome analysis is that some of the highly repetitive genes, including surface protein adhesins *ebpS* (elastin-binding) and *srdH* (unknown binding) (42), escaped ortholog clustering by reciprocal blast and were reported as distinct genes for each species. However, barring these few examples, most orthologs were successfully clustered together. Functional analysis of these genes showed that many of the species-restricted core genes encoded skin-relevant functions, potentially allowing species to co-colonize by using distinct molecular pathways. A large proportion (∼19%) of the species-restricted core genes were hypothetical, and future analysis of these genes could help identify novel functions unique to each species. For example, all *S. capitis* genomes carried a poorly characterized locus for gallidermin-type lantibiotic that might be crucial for competing with other skin resident microbes (43).

Finally, we used *in vitro* growth and RNA-Seq to characterize the inter-and intra-species phenotypic and transcriptional variation under similar growth conditions as a potential indicator of phylogenetic and ecological variation. Although the expression pattern of core genes was more similar between conspecific isolates, overall, wide variation was observed in gene expression between isolates highlighting the importance of gene regulation in amplifying the genetically encoded differences between each isolate.

Given the ease of culturing staphylococci, their survival outside the host environment, and our ability to genetically manipulate members of this genus (14), staphylococci hold great potential to provide strain-specific functions needed to maintain or restore a healthy skin barrier. For instance, the antibiotic-resistant pathogens such as methicillin-resistant *S. aureus* have reduced the efficacy of traditional antibiotics and requires innovative solutions to combat this crisis. Traditional antagonism assays have shown that staphylococci are a rich source of antimicrobial peptides including lantibiotics and nonribosomally synthesized antibiotics such as lugdunin that have been shown to target *S. aureus* (7, 8, 44, 45). A systematic genome mining approach could enable the discovery of additional novel and unique functions to restrict colonization by deadly pathogens. Our current study provides an important resource for such ambitious future studies.

## MATERIALS AND METHODS

### Healthy Volunteer recruitment and sampling

Healthy adult male and female volunteers 18–40 years of age were recruited from the Washington, DC metropolitan region. This natural history study was approved by the Institutional Review Board of the National Human Genome Research Institute (<u>clinicaltrials.gov/NCT00605878</u>) and the National Institute of Arthritis and Musculoskeletal and Skin Diseases (<u>clinicaltrials.gov/NCT02471352</u>) and all volunteers provided written informed consent prior to participation. Sampling was performed as described previously (23).

### 16S rRNA amplicon sequencing and analysis

Sample processing and sequencing were performed as described previously (23). Sequencing data was curated to include samples from fourteen distinct body sites collected from twenty-two healthy volunteers (HVs) (**Fig. S1**). 16S rRNA amplicon (V1-V3) sequencing data were processed using the DADA2 pipeline version v1.2.0 (46) to produce abundance estimates for error corrected amplicon sequence variants (ASVs). Reads were truncated to 375 nt with a max expected error (maxEE) of 2. The dada command (DADA2) was used to infer the composition of the sample with additional parameters specific to pyrosequencing data (HOMOPOLYMER_GAP_PENALTY=-1, BAND_SIZE=32). Chimeras were removed with the removeBimeraDenovo command. Taxonomic assignments were made using the DADA2 assignTaxonomy command with a curated RefSeq database (https://zenodo.org/record/3266798) and minimum bootstrap of 75. Taxonomic assignments were refined using BLAST to correct cases where a given ASV doesn’t uniquely identify a species (see below). While we have shown previously that human-associated staphylococci can be differentiated using V1-V3 sequence data of sufficient length (20), we nevertheless tested the accuracy of our species-level assignments using simulations. Briefly, 74 Type sequence *Staphylococcus* isolates from the Ribosomal Database Project (47) were used to generate simulated sequences with an error profile similar to the one observed for pyrosequencing: 0.5% base changes, 0.1% inserts, 0.1% deletions (n=10 per reference sequence). Simulated sequences were trimmed to a 375 base V1-V3 region analogous to the real data and classified. The following species could not be separated, due to lack of variation in the selected V1-V3 region: *S. capitis/S. caprae*, *S. agnetis/S.* hyicus, *S. pseudointermedius/S. intermedius/S. delphini*. In addition, *S. argenteus* isn’t distinguishable from *S. aureus* using V1-V3; however, these two species are in general known to be difficult to differentiate (48). In addition to these, we could not assign taxonomy to two prominent species numbered Species-4 and Species-45. These were labelled as *S. epidermidis* group and *S. hemolyticus* group, respectively, denoting their closest taxonomically related species. ASV abundance estimates were then merged with sample metadata into a phyloseq (49) object for further processing.

### Statistical analysis of 16S rRNA amplicon sequencing data

All analyses were performed in RStudio using the R v4.2.0 on the phyloseq object. For staphylococcal community analyses, the phyloseq object was trimmed to retain *Staphylococcus* ASVs using the subset_taxa function. ASVs that were present at ≥ 1% relative abundance in at least one sample were included. Dataset was further filtered to only retain ASVs present in more than three volunteers. This resulted in a final phyloseq object with 71 unique ASVs present in 298 samples. Read count for these 71 ASVs ranged from 99 to 176996. ASVs that belonged to the same species were merged using the tax_glom function within phyloseq, resulting in a total of 17 staphylococcal species. The distribution of these species was displayed using barplots, prevalence-abundance dotplots and boxplots. Non-parametric Wilcoxon rank-sum test was used to calculate statistical differences between microbial populations at different body-sites. P values below a standard alpha value (P value < 0.05) were considered significant. For beta-diversity measurement, phyloseq object with unmerged ASVs (before tax-glom) was used. Arcsine transformation was used for variance stabilization of OTU counts. Beta-diversity was visualized by Principal Coordinate Analysis (PCoA) of the Bray-Curtis dissimilarity metric generated using vegdist in the vegan v2.6-2 package. Effect size of individual ASVs was calculated with the envfit function (vegan). Variation by body-site was measured using PERMANOVA with the adonis function (vegan), using 10,000 permutations.

### Isolate collection

Skin and nasal cultures were obtained with Catch-all Collection Swabs (Epicentre) pre-moistened with Fastidious Broth (Remel), placed in 2.0ml Fastidious Broth supplemented with 10% glycerol, and frozen at −80°C. Swabs were thawed, vortexed, serial diluted, and plated on Tryptic Soy Agar with 5% Sheep Blood (Remel). After overnight incubation at 37°C, colonies were picked and stored in TSB with 20% glycerol. Colonies were screened by PCR for *S. capitis* using ScapF (5’-GCTAATTTAGATAGCGTACCTTCA −3’) and ScapR (5’- CAGATCCAAAGCGTGCA -3’) (50), *S. epidermidis* using Se705-1 (5’- ATCAAAAAGTTGGCGAACCTTTTCA -3’) and Se705-2 (5’- CAAAAGAGCGTGGAGAAAAGTATCA -3’) (50) and *S. hominis* using hom-F (5’- TACAGGGCCATTTAAAGACG - 3’) and hom-R (5’- GTTTCTGGTGTATCAACACC -3”) (51). Species taxonomy of isolates was confirmed by sequencing 16S rRNA gene using Sanger sequencing.

### Whole genome sequencing

Individual staphylococcal colonies were streaked on blood agar for two passages. Isolates were grown overnight in Tryptic Soy Broth at 37° C, pelleted with centrifugation, and genomic DNA was extracted using the Promega Maxwell Tissue DNA Kit with the addition of Readylyse Lysozyme Solution (Epicentre) and Lysostaphin (Sigma). DNA was treated with RNase, re-purified with the Genomic DNA Clean and Concentrator Kit (Zymo), and quantified using a Nanodrop spectrophotometer and Qubit (ThermoFisher). 1.0ng of bacterial DNA was used as input into the Nextera XT Sample prep kit (Illumina) as suggested by manufacturer. Nextera libraries were generated from the genomic DNA and sequenced using a paired-end 300-base dual index run on an Illumina MiSeq to generate 1 million to 2 million read pairs per library for ∼80x genome coverage. 79 *S. capitis*, 73 *S. epidermidis*, and 121 *S. hominis* isolates (273 total) from eighteen body sites of fourteen healthy volunteers were sequenced.

### Whole genome assembly

Nextera libraries for each isolate were multiplexed on a Novaseq 6000. Reads were subsampled to 80x coverage using seqtk (version 1.2), assembled with SPAdes (version 3.14.1) (52) and polished using bowtie2 (version 2.2.6) and Pilon (version 1.23) (53). To achieve full reference genomes for select isolates, genomic DNA was sequenced on the PacBio Sequel II platform (version 8M SMRTCells, Sequel II version 2.0 sequencing reagents, 15 hr movie collection). The subreads were assembled using Canu v2.1 and polished using the pb_resequencing workflow within PacBio SMRTLink v.9.0.0.92188. Genome annotation was performed using National Center for Biotechnology Information (NCBI) Prokaryotic Genome Annotation Pipeline (PGAP: https://www.ncbi.nlm.nih.gov/genome/annotation_prok/).

### Genome quality control

Genomes with ≥ 95% fastANI distances were considered to belong to the same species (25). All 273 genomes were quality filtered according to completeness (≥ 98%) and contamination (≤ 5%) using CheckM (54). dRep v3.2.2 (55) was used to de-replicate and collapse highly similar genomes using a >99.9% ANImf threshold and a minimum alignment coverage of 50%, as used previously (56). A representative genome from each clonal cluster was chosen for pan-genome analysis based on dRep’s score-based system, which incorporates completeness, contamination, strain heterogeneity, N50, genome length and centrality. The following command was used: *dRep dereplicate -p 10 -comp 98 -con 5 -sa 0.999 -nc 0.5 -g *.fasta*

### Species-level pan-genome

Dereplicated draft genomes, including plasmids, were annotated using prokka v1.14.6 (57). A customized protein database was generated for prokka annotation using the nine staphylococcal reference genomes sequenced in this study using either the PacBio or Oxford Nanopore technology. This was to ensure consistent annotation across all genomes. Prokka generated *gff3 output was passed on to panaroo v3.1.2 (58) for species-level pan-genome analysis. Panaroo was run under the strict mode without merging paralogs. The initial sequence identity threshold was 98% with subsequent clustering into protein families being performed using a threshold of 70% identity. Core genes were defined using a 99% presence threshold. A core gene alignment was generated using the default mafft aligner and the -a flag, which was used for generating phylogenetic trees. Panaroo command: *panaroo -i *.gff -o results -clean-mode strict -refind_prop_match 0.7 -alignment core -codons -t 24*

### Genus-level pan-genome

We employed a reciprocal-best-blast approach to cluster the species-level pan-genome references (containing a representative sequence from each orthologous gene cluster detected in the pan-genome) at the protein level (≥ 40% identity and ≥ 50% coverage). This resulted in clustering of 78-93% of core genes from each species-level pan-genome to provide a merged genus pan-genome.

### Pan-genome analyses

Gene presence absence matrices from Panaroo and three-way-reciprocal blast were used for individual species and genus pan-genome analyses, respectively, using R v4.2.0. micropan package v2.1 (59) was run with default parameters for principal component analysis (PCA). Cluster analysis to identify genome grouping based on gene content was performed using dist() and hclust functions in base R. pheatmap v1.0.12 was used for generating heatmaps. Venneuler 1.1.3 was used to make proportional Venn diagrams. Tanglegram showing phylogeny versus pan-genome tree was generated using dendextend package v1.16.

### Functional annotation and reconstruction of metabolic pathways

Prokka provided default annotations for most genes. In addition to this, thorough functional annotations and pfam domain predictions were carried out by eggNOG-mapper v2 (web version) using eggNOG 5.0 database, DIAMOND aligner, and the following flags “–evalue 0.001 –score 60 –pident 40 –query_cover 20 –subject_cover 20 –itype CDS –translate –tax_scope auto –target_orthologs all –go_evidence all –pfam_realign denovo”. Panaroo output file pan_genome_reference.fa, containing reference sequences of all the genes found in a species pan-genome, was used as an input for annotation.

Completeness of Kyoto Encyclopedia of Genes and Genomes (KEGG) pathways was tested using a previously described pipeline (60). Bacterial pathways that showed ³ 75% completeness in at least one genome were kept for downstream analyses.

Functional enrichment analysis of KEGG pathways was performed using MicrobiomeProfiler, a R-shiny package based on clusterProfiler (61). KO identifiers assigned by eggNOG-mapper v2 were used as input for the analysis.

### Phylogenetic tree construction

Phylogenetic tree for each species was constructed using the core-gene alignment generated by panaroo. ModelFinder (62) implemented in IQ-TREE 2 (63) was employed to find the best substitution and site heterogeneity models. GTR+F+I+G4, GTR+F+R2 and GTR+F+R5 models were selected for *S. capitis*, *S. epidermidis*, and *S. hominis*, respectively to construct maximum likelihood (ML) trees using RAxML-NG v1.1.0 (64) with 1,000 bootstrap replicates. Each species tree was rooted using genomes from the other two species as an outgroup. For this rooting, a genus-level phylogenetic tree was built using GTR+F+R5 model on the core-gene alignment of 665 genus-core genes generated by panaroo using all 126 genomes and the same threshold values as outlined above. iTOL v6 (65) was used for tree display and annotation. RAxML-NG command: *raxml-ng -all -msa core_gene_alignment_filtered.aln -model -prefix -seed -threads -bs-metric fbp,tbe -bs-trees 1000*

### pan-GWAS analysis

Scoary v1.6.16 (66) was used to identify overrepresentation of genes within each phylogenetic clade for individual species. The input trait file consisted of dummy variables (0 and 1) to denote clade A and clade B membership. gene_presence_absence_roary.csv output from panaroo was used for gene content differences between species genomes. Scoary was run for each species using the following command: *scoary -g gene_presence_absence_roary.csv -t traits_clades.csv -no_pairwise -o results p 0.05 -c BH --threads 10*

### Isolate selection for growth curves

A greedy algorithm was used to choose a subset of staphylococcal isolates for growth curve analysis. Briefly, *gene_presence_absence.Rtab* output file from panaroo was used to iteratively select genomes that best improve the representation of the species-level pan-genome. Initially, genomes were selected to cover the core+softcore+shell genes (genes found in at least 15% of genomes) using the minimum number of genomes. Then additional isolate genomes were added to increase the coverage of cloud genes (present in <15% of genomes) such that > 88% of the pan-genome (without singletons) was encoded by final isolate genomes. In some cases, we forced the inclusion of genomes that were phenotypically valuable or had finished genomes. The final dataset included 15 *S. epidermidis* isolates (75.4% total pan-genome; 96.1% without singletons), 12 *S. capitis* (94.1%; 99.6%), and 14 *S. hominis* isolates (68.2%; 88.7%).

### Growth analysis

All isolates were cultivated overnight, at 37 °C aerobically, in Brain Heart Infusion with 0.5% Yeast extract broth (BHI-YE) from frozen glycerol stocks. Overnight cultures were used as inocula (1:100) for BHI-YE and artificial skin media ES and ESL (Pickering Laboratories Inc Catalog #s 1700-0023 and 1700-0556 respectively), which were distributed in 96-microwell Nunc™ Edge plate (Catalog # 267544) (200ul/well in duplicate). Plates were sealed using Parafilm™ to reduce evaporation and incubated in BioTek Epoch 2 microplate reader at 34° C with continuous shaking at 180 rpm. Kinetic growth data was monitored by OD measurement at 600 nm every 30 minutes for up to 24 hours. A minimum of 4 biological replicates were generated for each isolate-medium pair by repeating the growth curves on separate days. Biological replicate data was pooled and area under the curve (AUC) quantified using Growthcurver (67). Data analyses and visualization was carried out using the R software. Composition of ES and ESL media are given in **Table S8.**

### RNA extraction and library preparation

Isolates for RNA-Seq analysis were grown following the same protocol as for growth curve analysis, except for using an Eppendorf deep 96-wells plate (2 ml) to accommodate larger culture volumes necessary to harvest sufficient RNA for sequencing. Cultures were grown to mid-log phase, and up to 6 ml of culture per isolate per medium was harvested using Bacterial RNAprotect (Qiagen 76506). Three biological replicates were generated for each condition.

For RNA extraction we used ZymoBiomcs DNA/RNA Miniprep Kit (R2002) to isolate bacterial RNA by adding 600μl RNA Shield to resuspend the bacterial pellets. After following all the steps outlined in the kit catalog, final total RNA was eluted in with 50μl nuclease free water. An extra DNase step was added to ensure complete removal of genomic DNA contamination. This involved digestion with 1μl of rDNase (Thermo Fisher AM2222) at 37°C for 30 min, followed by inactivation with 0.1 volume of DNase inactivation reagent. The extracted total RNA was cleaned up using NEB’s Monarch RNA Cleaning kit (T2030L), then eluted with 10-20μl nuclease-free water. RNA was quantified by Nanodrop and Qubit and the quality check was performed using bioanalyzer. Ribosomal RNA was removed by Qiagen’s FastSelect - 5S/16S/23S (335927). RNA fragmentation was set at 89°C for varying times depending on sample RIN (RNA integrity number), then gradually cooled using a gradient from 75°C to 25°C. After cleanup using QIAseq, Illumina stranded total RNA prep kit (20040529) was used to synthesize first and second strand cDNA. Agencourt AMPure XP beads (A63881) were used for cDNA cleanup. Adenine nucleotide was added to the 3’ ends for ligating RNA-index anchors to the double-stranded cDNA fragments, followed by another cleanup using AMPureXP beads. Selective PCR amplification of the anchor-ligated DNA fragments was done using unique-dual Index (20627581) to generate an RNA-Seq library. Libraries were assayed for quality and concentration by Agilent 2100 Bioanalyzer and DNA 1000 kit, then sent to NISC for sequencing.

### Bioinformatic analysis of RNA-seq data

Libraries were sequenced on the Illumina NovaSeq 6000 platform at the NIH Intramural Sequencing Center to a target depth of 50 million 2×150 paired-end reads per sample. Reads were trimmed for adapters with cutadapt v3.4 using the parameters “--nextseq-trim 20 -e 0.15 - m 50” (https://journal.embnet.org/index.php/embnetjournal/article/view/200) and checked for quality with PRINSEQ-lite v0.20.4 using the parameters “-lc_method entropy -lc_threshold 70 - min_len 50 -min_qual_mean 20 -ns_max_n 5 -min_gc 10 -max_gc 90” (68). Reads with less than 50 bp length after trimming were removed.

Rockhopper v 2.0.3 (69) was used to align reads from each sample to a reference genome. Each isolate had its own reference genome to ensure alignment of non-core gene reads. Raw read counts per gene were obtained from rockhopper, and those mapping to ncRNAs, rRNAs and tRNAs were removed from the analysis. Initial analysis was performed using a subset of 1647 genus-core genes that had at least one read in all samples. For this purpose raw reads were normalized using the estimateSizeFactors() function and transformed using the varianceStabilizingTransformation() function in the DESeq2 package (70) prior to analysis.

Principal components were determined using the prcomp() function. DESeq2 was used to calculate differentially expressed genes (log_2_ fold change ≥ 1 or ≤ −1 and adjusted P value < 0.05) in ES medium relative to BHI-YE in individual isolates to measure the contribution of all the genes encoded within a genome. Differential expression was determined using ’apeglm’ for LFC shrinkage (71). Heatmaps were generated using pheatmap and ComplexHeatmap R packages.

## Data Availability

Genome data are deposited under the NCBI BioProjects PRJNA694925 and PRJNA986048. Some amplicon data were published previously (N = 145; PRJNA46333) (72) and the remainder are new to this study (N = 168; PRJNA46333). RNAseq data have been deposited under BioProject PRJNA694925. Code for this project along with the phyloseq object are available at https://github.com/skinmicrobiome/Joglekar_Staphylococcus_2023

## ACKNOWLEDGEMENTS

We thank Lukian Robert and Qiong Chen for assisting with culturing, DNA extraction, and library preparation. The computational resources of the NIH High-Performance Computation Biowulf Cluster (http://hpc.nih.gov) were used for this study. This work was supported by the Intramural Research Programs of NHGRI and NIAMS.

## AUTHOR CONTRIBUTIONS

P.J., S.C., J.A.S., and H.H.K. conceived and designed the study. P.J., and S.C. performed amplicon and pan-genome analysis. P.J. performed growth curves and RNA-Seq analysis. H.H.K. collected samples. P.J., and C.D. cultured bacteria and C.D. prepared DNA sequencing libraries; S-Q.L-L. extracted RNA and prepared RNA sequencing libraries. NISC performed sequencing. S.C. managed and preprocessed data. S.C. and S.S.K. provided software and bioinformatic support. P.J., S.C., J.A.S., wrote the manuscript. J.A.S. and H.H.K. provided funding. All authors reviewed and approved the manuscript.

## DECLARATION OF INTERESTS

The authors declare no competing interests.

## Notes

### Competing Interest Statement

The authors have declared no competing interest.

