## Supplementary Figures and Tables for "Integrated genomic and functional analyses of human skin-associated *Staphylococcus* reveals extensive inter- and intra-species diversity"

**Julia A. Segre**

Building 49, Room 4a26,

49 Convent Dr. 4442,

Bethesda, MD 20892-4442

(301) 402-2314

**This PDF file includes:**

- Figures S1 to S10
- Tables S1, S5, S8 and S9
- Legends for Datasets S2, S3, S4, S6, S7, S10
- SI References

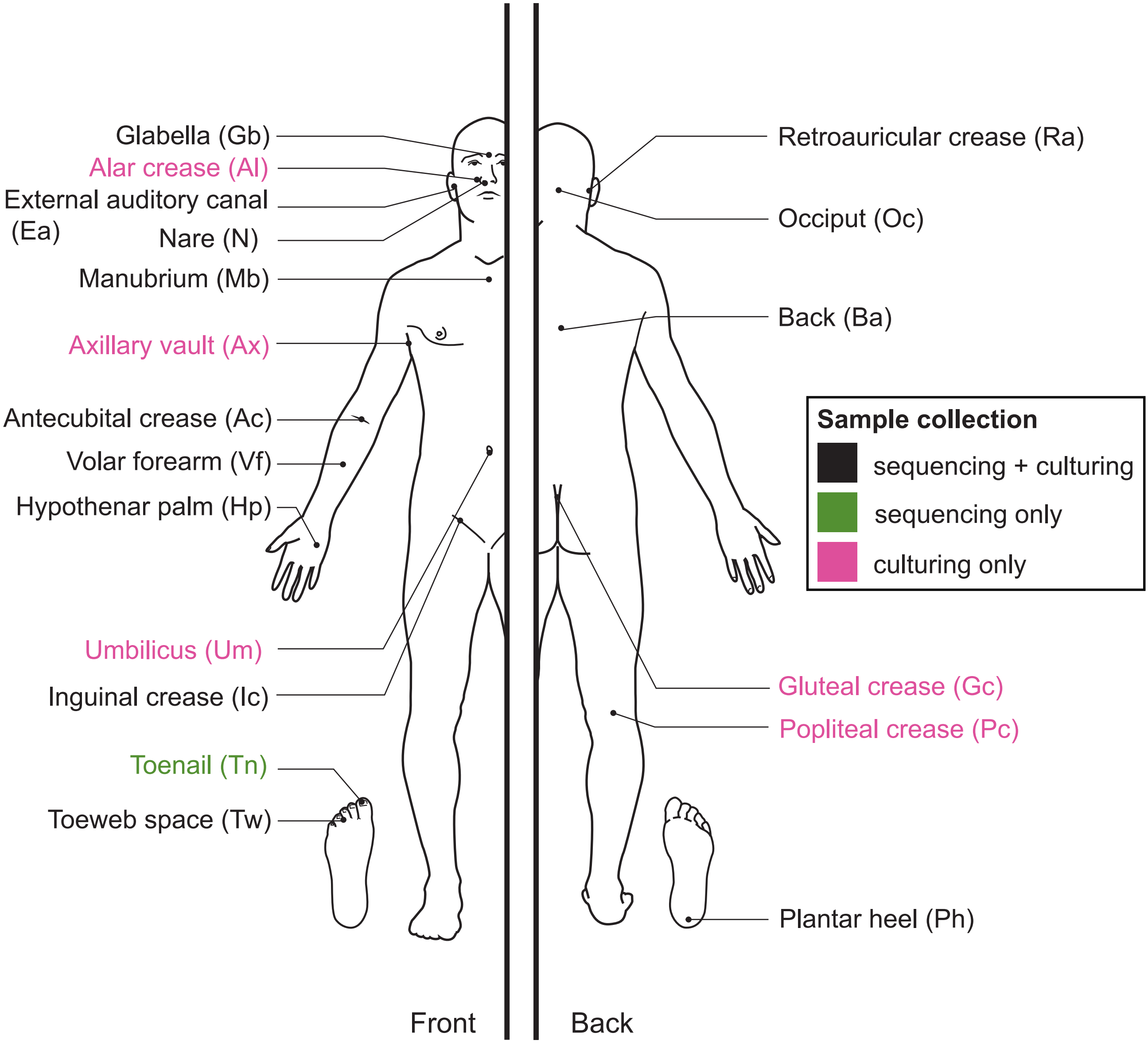
Glabella (Gb)
Alar crease (Al)
External auditory canal (Ea)
Nare (N)
Manubrium (Mb)
Axillary vault (Ax)
Antecubital crease (Ac)
Volar forearm (Vf)
Hypothenar palm (Hp)
Umbilicus (Um)
Inguinal crease (Ic)
Toenail (Tn)
Toeweb space (Tw)
Retroauricular crease (Ra)
Occiput (Oc)
Back (Ba)
Sample collection
sequencing + culturing
sequencing only
culturing only
Gluteal crease (Gc)
Popliteal crease (Pc)
Plantar heel (Ph)
Front
Back

**Figure S1**. **Body sites used for collecting samples from healthy volunteers for 16S rRNA amplicon sequencing and culturing of staphylococcal isolates**.

Sites shown in black represent those for which both types of samples were collected. Toenail samples (green) were only collected for sequencing. Umbilicus, Gluteal crease, and Popliteal crease samples (all pink) were collected for culturing alone. Abbreviation shown in a bracket next to each body site were used to denote the site throughout the manuscript.

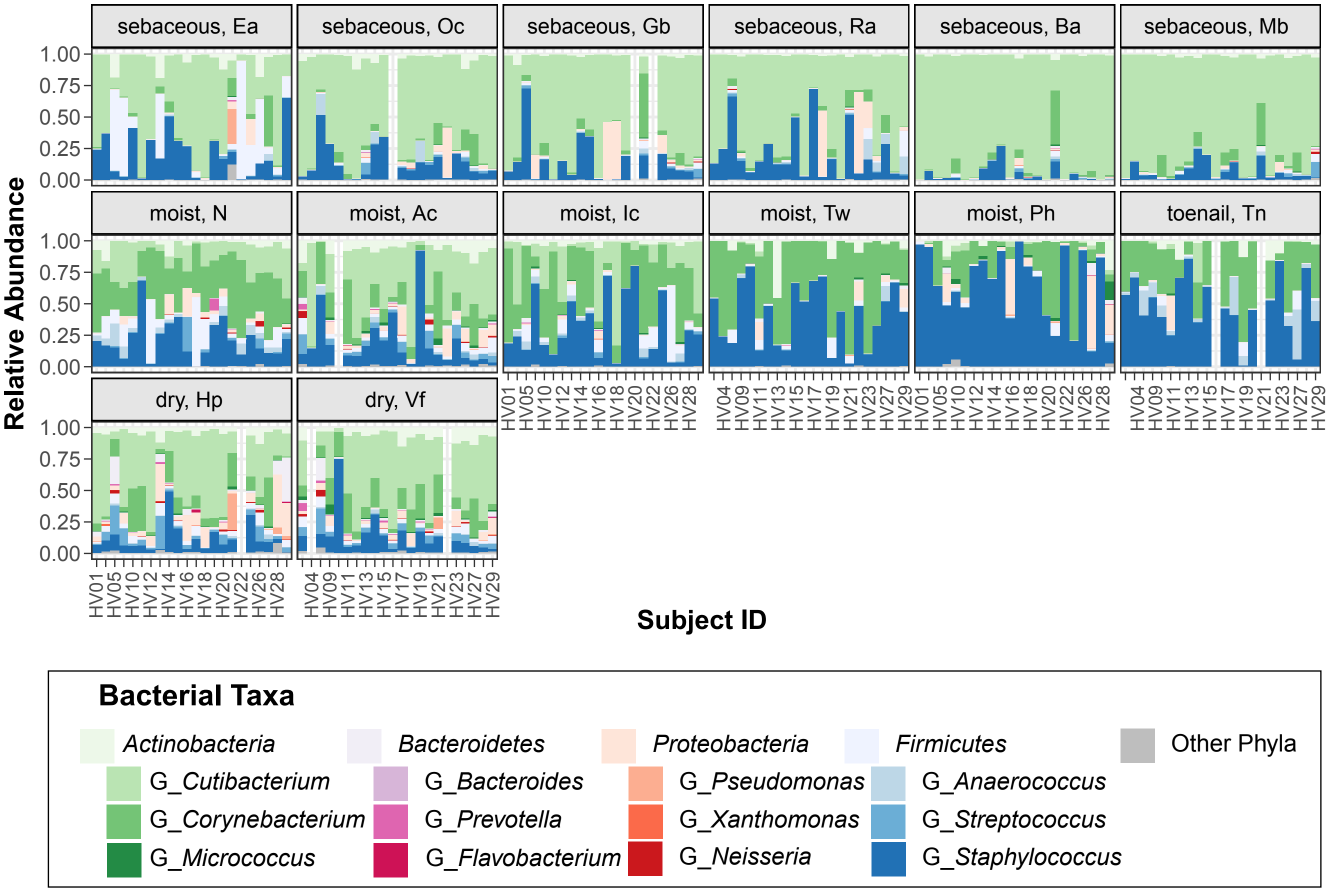
sebaceous, Ea
sebaceous, Oc
sebaceous, Gb
sebaceous, Ra
sebaceous, Ba
sebaceous, Mb
moist, N
moist, Ac
moist, Ic
moist, Tw
moist, Ph
toenail, Tn
dry, Hp
dry, Vf
Relative Abundance
1.00
0.75
0.50
0.25
0.00
HV01 HV05 HV10 HV12 HV14 HV16 HV18 HV20 HV22 HV26 HV28
HV04 HV09 HV11 HV13 HV15 HV17 HV19 HV21 HV23 HV27 HV29
Subject ID
Bacterial Taxa
Actinobacteria
Bacteroidetes
Proteobacteria
Firmicutes
Other Phyla
G_Cutibacterium
G_Corynebacterium
G_Micrococcus
G_Bacteroides
G_Prevotella
G_Flavobacterium
G_Pseudomonas
G_Xanthomonas
G_Neisseria
G_Anaerococcus
G_Streptococcus
G_Staphylococcus

**Figure S2**. **Bacterial diversity on healthy human skin represented by major phyla and genera colonizing distinct body sites.**

Barplots display the relative abundance of major bacterial taxa at various body sites as displayed by facets. Colors represent taxa as shown in the accompanying legend. Each bar represents one volunteer. Empty bars represent missing data. Refer to Fig. S1 for body site abbreviation.

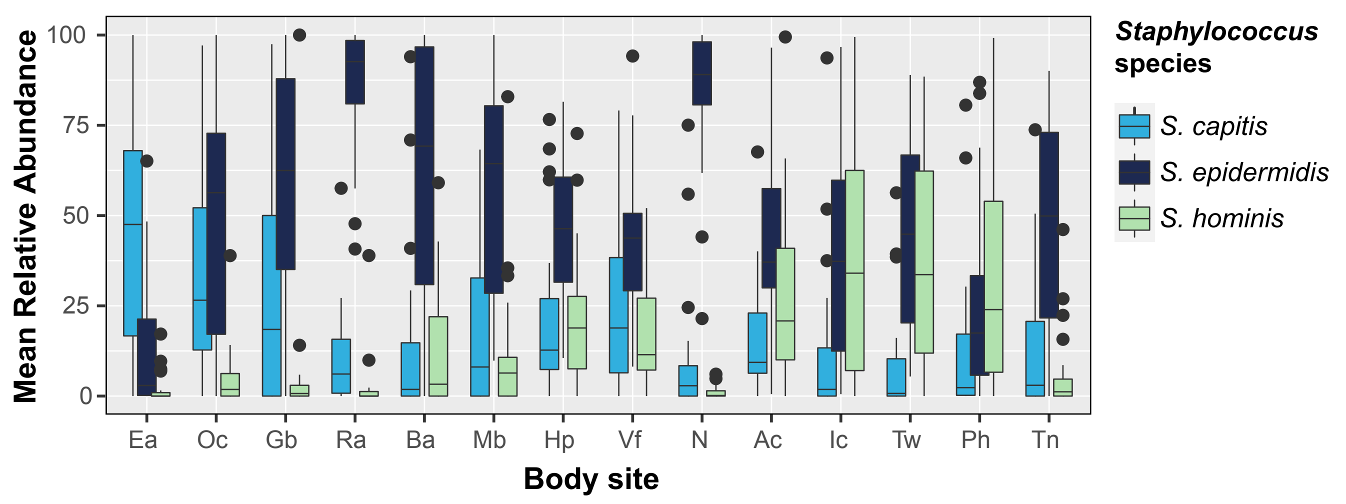

Mean Relative Abundance
100
75
50
25
0
Ea
Oc
Gb
Ra
Ba
Mb
Hp
Vf
N
Ac
Ic
Tw
Ph
Tn
Body site
Staphylococcus species
S. capitis
S. epidermidis
S. hominis

**Figure S3. Mean relative abundance (MRA) of the three most prominent species within staphylococcal communities**

Boxplots represents the MRA of the three staphylococcal species across all body sites as shown on the x-axis. Boxplot colors represent individual species. The center black line within each boxplot represents the median value, with edges showing the first and third quartiles. Refer to Fig. S1 for body site abbreviation.

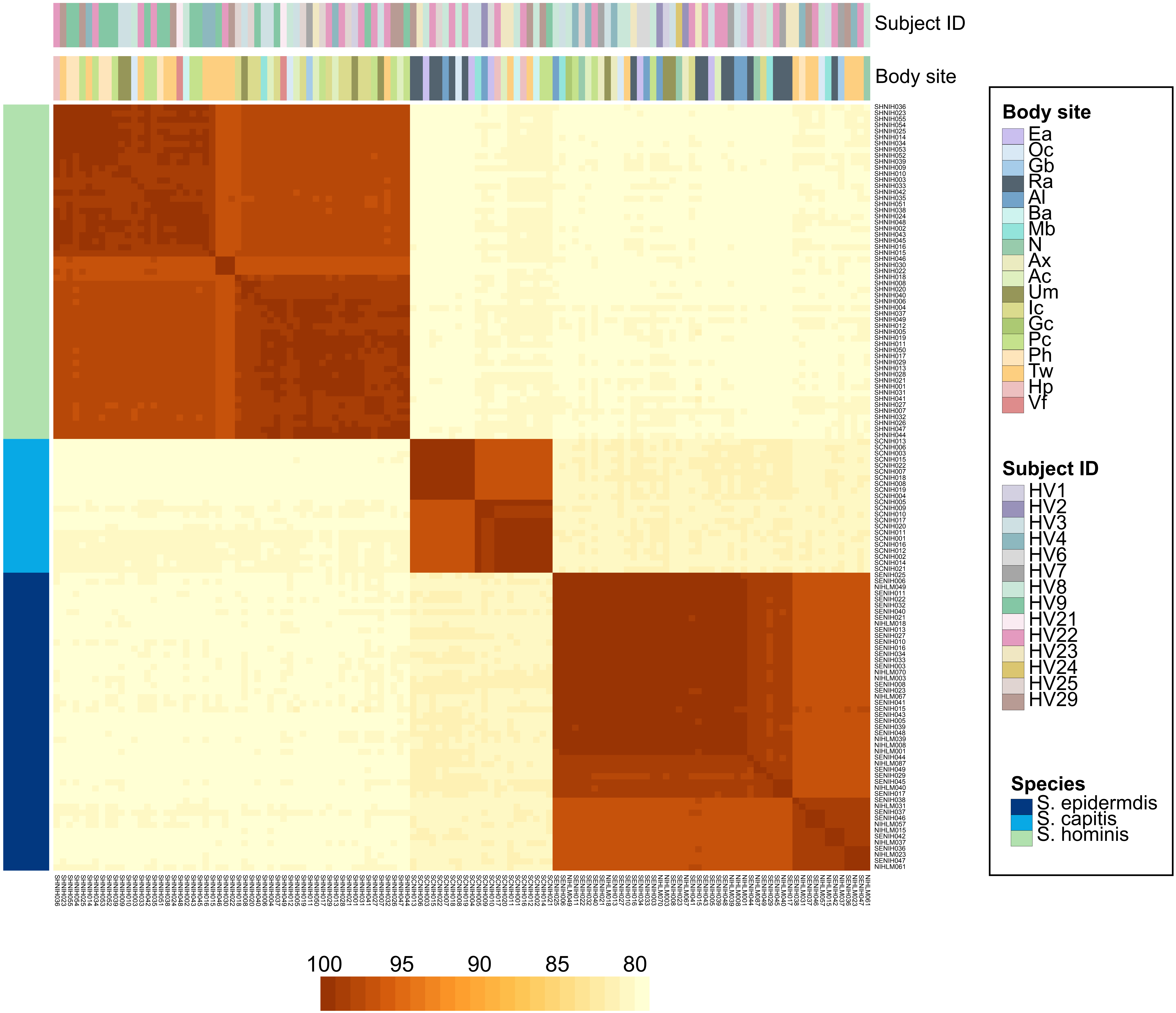
Subject ID
Body site
Body site
Ea
Oc
Gb
Ra
Al
Ba
Mb
N
Ax
Ac
Um
Ic
Gc
Pc
Ph
Tw
Hp
Vf
Subject ID
HV1
HV2
HV3
HV4
HV6
HV7
HV8
HV9
HV21
HV22
HV23
HV24
HV25
HV29
Species
S. epidermdis
S. capitis
S. hominis
100
95
90
85
80

**Figure S4**. **Heat-map of pairwise fastANI comparison of de-replicated, non-clonal genomes of *S. epidermidis, S. capitis*, and *S. hominis*.**

Colored bars on the top denote the healthy volunteer and body site of isolation of each labeled genome. Species are denoted by the side bar on the left. Refer to Fig. S1 for body site details.

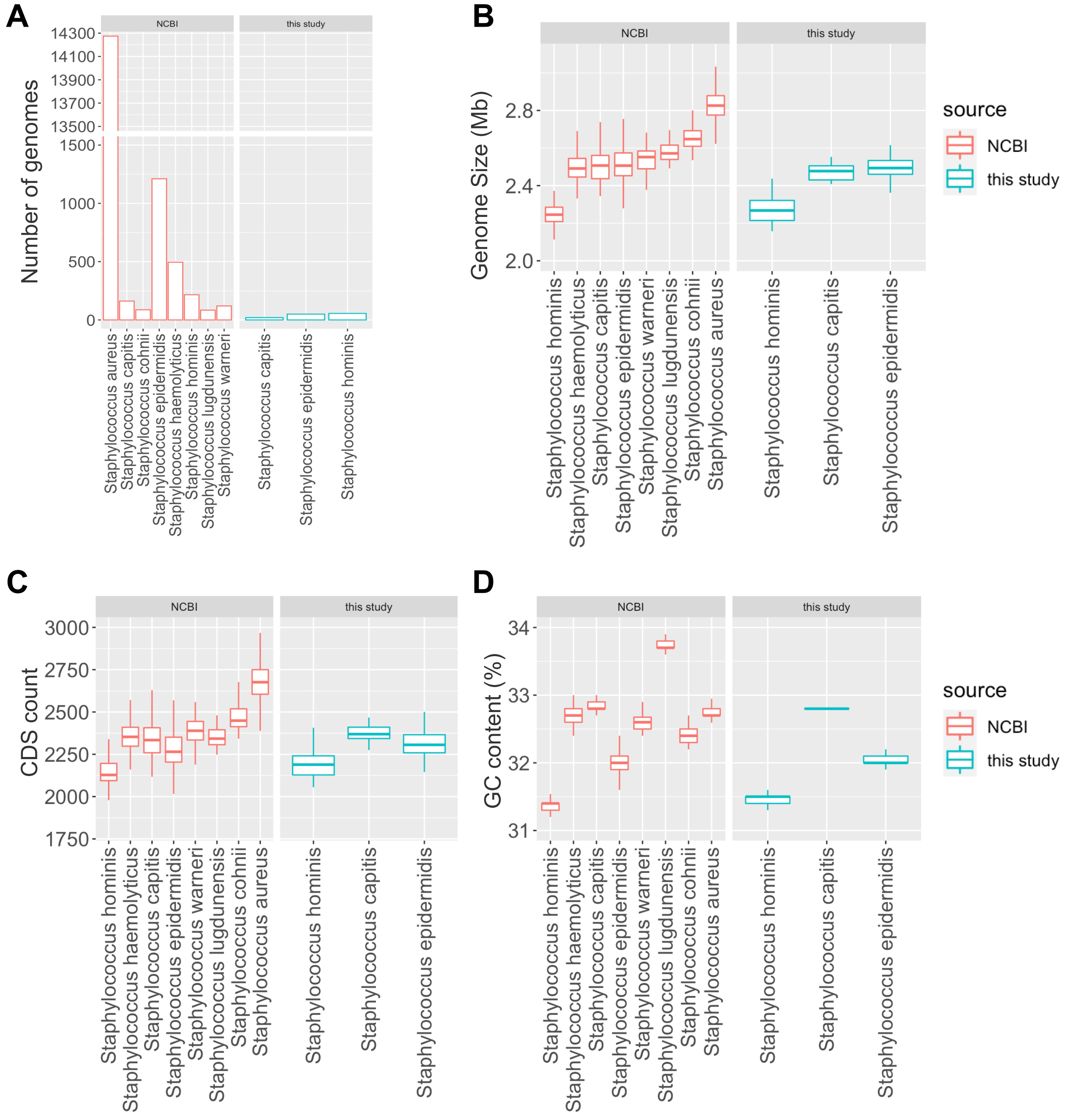
A
NCBI
this study
Number of genomes
14300
14100
13900
13700
13500
1500
1000
500
0
Staphylococcus aureus
Staphylococcus capitis
Staphylococcus cohnii
Staphylococcus epidermidis
Staphylococcus haemolyticus
Staphylococcus hominis
Staphylococcus lugdunensis
Staphylococcus warneri
Staphylococcus capitis
Staphylococcus epidermidis
Staphylococcus hominis
B
NCBI
this study
Genome Size (Mb)
2.8
2.4
2.0
Staphylococcus hominis
Staphylococcus haemolyticus
Staphylococcus capitis
Staphylococcus epidermidis
Staphylococcus warneri
Staphylococcus lugdunensis
Staphylococcus cohnii
Staphylococcus aureus
Staphylococcus hominis
Staphylococcus capitis
Staphylococcus epidermidis
source
NCBI
this study
C
NCBI
this study
CDS count
3000
2750
2500
2250
2000
1750
Staphylococcus hominis
Staphylococcus haemolyticus
Staphylococcus capitis
Staphylococcus epidermidis
Staphylococcus warneri
Staphylococcus lugdunensis
Staphylococcus cohnii
Staphylococcus aureus
Staphylococcus hominis
Staphylococcus capitis
Staphylococcus epidermidis
D
NCBI
this study
GC content (%)
34
33
32
31
Staphylococcus hominis
Staphylococcus haemolyticus
Staphylococcus capitis
Staphylococcus epidermidis
Staphylococcus warneri
Staphylococcus lugdunensis
Staphylococcus cohnii
Staphylococcus aureus
Staphylococcus hominis
Staphylococcus capitis
Staphylococcus epidermidis
source
NCBI
this study

**Figure S5**. **Boxplot representing genome characteristics of isolates used in the current study and comparing them to species-specific genomes present in NCBI.**

**(A)** Number of genomes of different species that were either present in the NCBI or were sequenced for this study. **(B)** Average genome sizes of each species. **(C)** Average number of protein-coding genes (CDS) that were present in the genome of each species. **(D)** Percent GC content of each species.

The center line within each boxplot represents the median value, with edges showing the first and third quartiles.

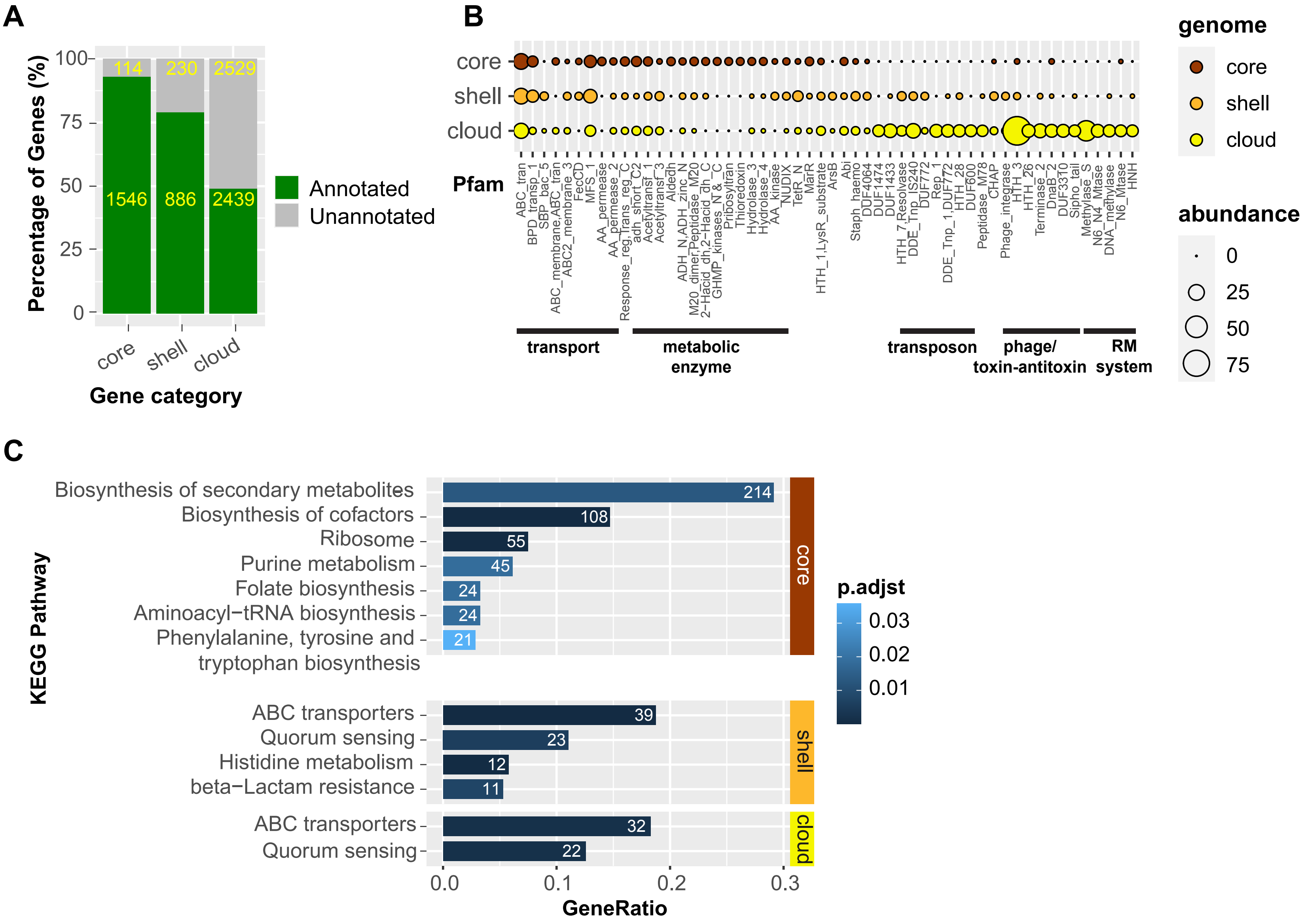
A
Percentage of Genes (%)
100
75
50
25
0
114
230
2529
1546
886
2439
core
shell
cloud
Gene category
Annotated
Unannotated
B
core
shell
cloud
Pfam
ABC_tran
BPD_transp_1
SBP_bac_5
ABC_membrane,ABC_tran
ABC2_membrane_3
FecCD
MFS_1
AA_permease
AA_permease_2
Response_reg,Trans_reg_C
adh_short_C2
Acetyltransf_1
Acetyltransf_3
Aldedh
ADH_N,ADH_zinc_N
M20_dimer,Peptidase_M20
2-Hacid_dh,2-Hacid_dh_C
GHMP_kinases_N,&_C
Pribosyltran
Thioredoxin
Hydrolase_3
Hydrolase_4
AA_kinase
NUDIX
TetR_N
MarR
HTH_1,LysR_substrate
ArsB
Abi
Staph_haemo
DUF4064
DUF1474
DUF1433
HTH_7,Resolvase
DDE_Tnp_IS240
DUF772
Rep_1
DDE_Tnp_1,DUF772
HTH_28
DUF600
Peptidase_M78
CHAP
Phage_integrase
HTH_3
HTH_26
Terminase_2
DnaB_2
DUF3310
Sipho_tail
Methylase_S
N6_N4_Mtase
DNA_methylase
N6_Mtase
HNH
transport
metabolic enzyme
transposon
phage/ toxin-antitoxin
RM system
genome
core
shell
cloud
abundance
0
25
50
75
C
KEGG Pathway
Biosynthesis of secondary metabolites
Biosynthesis of cofactors
Ribosome
Purine metabolism
Folate biosynthesis
Aminoacyl−tRNA biosynthesis
Phenylalanine, tyrosine and tryptophan biosynthesis
ABC transporters
Quorum sensing
Histidine metabolism
beta−Lactam resistance
ABC transporters
Quorum sensing
214
108
55
45
24
24
21
39
23
12
11
32
22
core
shell
cloud
p.adjst
0.03
0.02
0.01
0.0
0.1
0.2
0.3
GeneRatio

**Figure S6**. **Functional annotation of genus pan-genome.**

**(A)** Percentage of annotated genes shown by genus pan-genomic category. Actual number of genes in each group are shown in yellow inside each bar. **(B)** Combined data showing top 20 most represented Pfam domains in each pan-genomic category. Size of the bubbles represents the actual number of genes carrying the Pfam domain annotation within each category. **(C)** KEGG pathway enrichment analysis of genes in each category using KEGG KO identifiers.

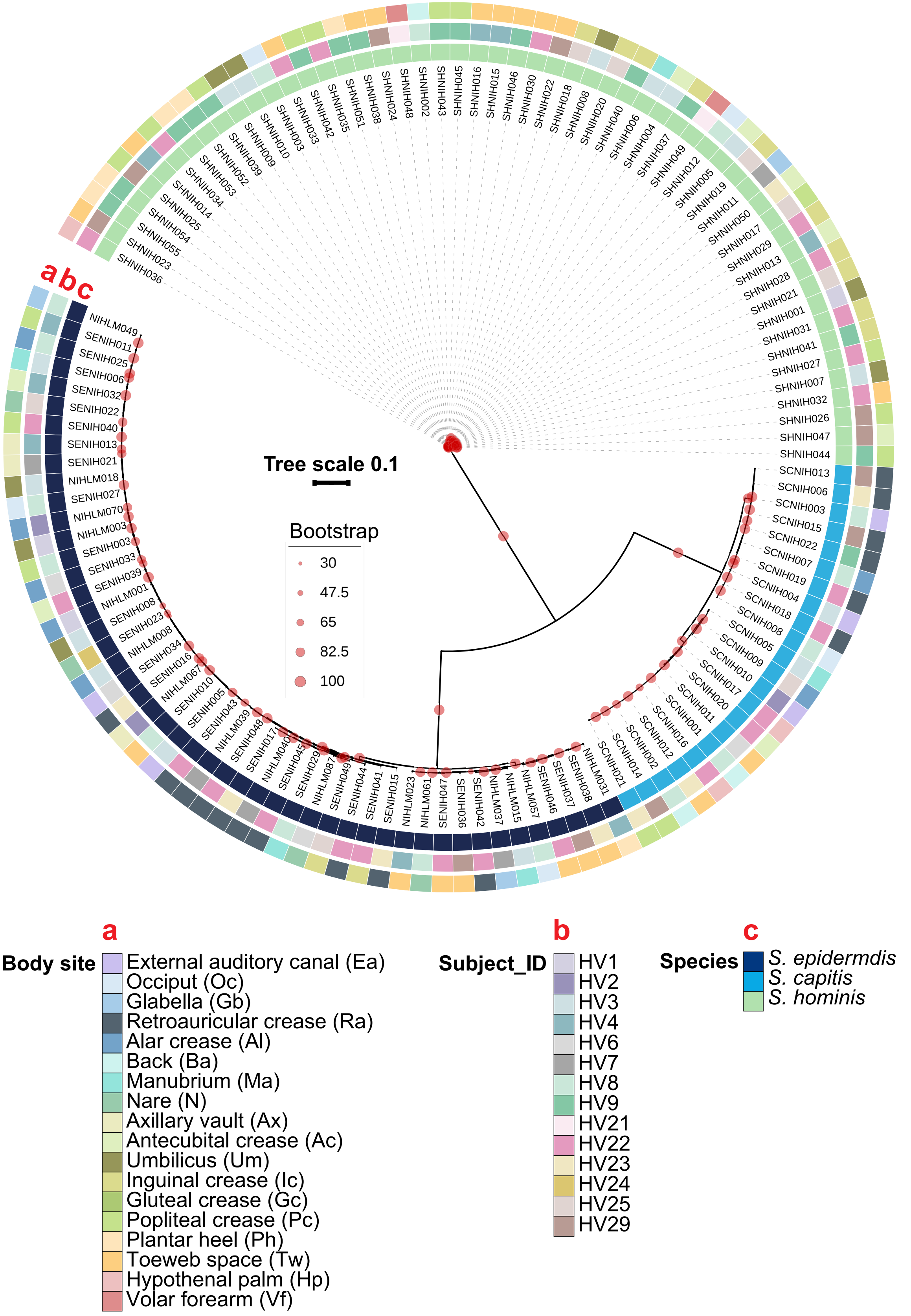
a b c
Tree scale 0.1
Bootstrap
30
47.5
65
82.5
100
SHNIH036
SHNIH023
SHNIH055
SHNIH054
SHNIH025
SHNIH014
SHNIH034
SHNIH053
SHNIH052
SHNIH039
SHNIH009
SHNIH010
SHNIH003
SHNIH033
SHNIH042
SHNIH035
SHNIH051
SHNIH038
SHNIH024
SHNIH048
SHNIH002
SHNIH043
SHNIH045
SHNIH016
SHNIH015
SHNIH046
SHNIH030
SHNIH022
SHNIH018
SHNIH008
SHNIH020
SHNIH040
SHNIH006
SHNIH004
SHNIH037
SHNIH049
SHNIH012
SHNIH005
SHNIH019
SHNIH011
SHNIH050
SHNIH017
SHNIH029
SHNIH013
SHNIH028
SHNIH021
SHNIH001
SHNIH031
SHNIH041
SHNIH027
SHNIH007
SHNIH032
SHNIH026
SHNIH047
SHNIH044
SCNIH013
SCNIH006
SCNIH003
SCNIH015
SCNIH022
SCNIH007
SCNIH019
SCNIH004
SCNIH018
SCNIH008
SCNIH005
SCNIH009
SCNIH010
SCNIH017
SCNIH020
SCNIH011
SCNIH001
SCNIH016
SCNIH012
SCNIH002
SCNIH014
SCNIH021
NIHLM031
SENIH038
SENIH037
SENIH046
NIHLM057
NIHLM015
NIHLM037
SENIH042
SENIH036
SENIH047
NIHLM061
NIHLM023
SENIH015
SENIH041
SENIH044
SENIH049
NIHLM087
SENIH029
SENIH045
NIHLM040
SENIH017
SENIH048
NIHLM039
SENIH043
SENIH005
SENIH010
NIHLM067
SENIH016
SENIH034
NIHLM008
SENIH023
SENIH008
NIHLM001
SENIH039
SENIH033
SENIH003
NIHLM003
NIHLM070
SENIH027
NIHLM018
SENIH021
SENIH013
SENIH040
SENIH022
SENIH032
SENIH006
SENIH025
SENIH011
NIHLM049
a
Body site
External auditory canal (Ea)
Occiput (Oc)
Glabella (Gb)
Retroauricular crease (Ra)
Alar crease (Al)
Back (Ba)
Manubrium (Ma)
Nare (N)
Axillary vault (Ax)
Antecubital crease (Ac)
Umbilicus (Um)
Inguinal crease (Ic)
Gluteal crease (Gc)
Popliteal crease (Pc)
Plantar heel (Ph)
Toeweb space (Tw)
Hypothenal palm (Hp)
Volar forearm (Vf)
b
Subject_ID
HV1
HV2
HV3
HV4
HV6
HV7
HV8
HV9
HV21
HV22
HV23
HV24
HV25
HV29
c
Species
S. epidermdis
S. capitis
S. hominis

**Figure S7**. **Phylogenetic tree of all 126 staphylococcal genomes based on the polymorphic sites detected in sequence alignment of select core genes (N = 665).** Bootstrap values of at least 80 percent are shown with red dots. For each species-level phylogenetic tree shown in Figure 5, the other two species served as an outgroup for rooting the tree based on this genus-level tree. Body site, healthy volunteer, and species of each genome are shown as sidebars. Refer to Fig. S1 for body site details.

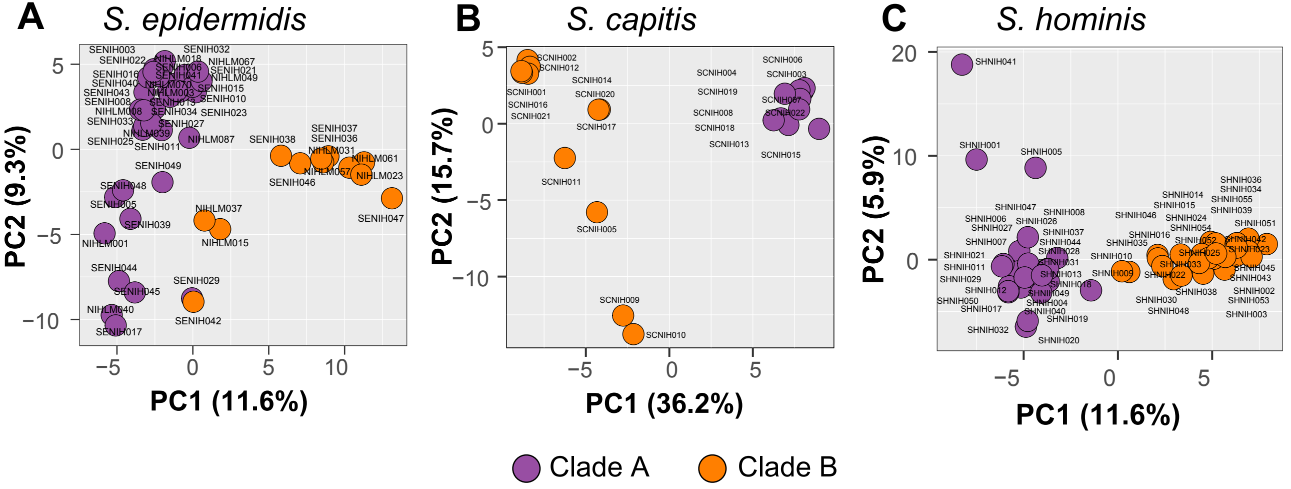
A
S. epidermidis
PC1 (11.6%)
PC2 (9.3%)
SENIH003
SENIH032
SENIH022
NIHLM018
NIHLM067
SENIH016
SENIH006
SENIH021
SENIH041
NIHLM049
SENIH040
NIHLM070
SENIH015
SENIH043
NIHLM003
SENIH010
SENIH008
SENIH013
NIHLM008
SENIH034
SENIH023
SENIH033
SENIH027
NIHLM039
SENIH025
SENIH011
NIHLM087
SENIH038
SENIH037
SENIH036
NIHLM031
NIHLM061
NIHLM057
NIHLM023
SENIH046
SENIH049
SENIH048
SENIH005
SENIH039
NIHLM001
NIHLM037
NIHLM015
SENIH047
SENIH044
SENIH045
SENIH029
NIHLM040
SENIH042
SENIH017
B
S. capitis
PC1 (36.2%)
PC2 (15.7%)
SCNIH002
SCNIH012
SCNIH001
SCNIH016
SCNIH021
SCNIH014
SCNIH020
SCNIH017
SCNIH011
SCNIH005
SCNIH009
SCNIH010
SCNIH004
SCNIH019
SCNIH008
SCNIH018
SCNIH013
SCNIH006
SCNIH003
SCNIH007
SCNIH022
SCNIH015
C
S. hominis
PC1 (11.6%)
PC2 (5.9%)
SHNIH041
SHNIH001
SHNIH005
SHNIH047
SHNIH008
SHNIH006
SHNIH026
SHNIH027
SHNIH037
SHNIH007
SHNIH044
SHNIH028
SHNIH021
SHNIH031
SHNIH011
SHNIH013
SHNIH029
SHNIH018
SHNIH012
SHNIH049
SHNIH050
SHNIH004
SHNIH017
SHNIH040
SHNIH019
SHNIH032
SHNIH020
SHNIH014
SHNIH015
SHNIH046
SHNIH024
SHNIH054
SHNIH016
SHNIH035
SHNIH052
SHNIH010
SHNIH025
SHNIH033
SHNIH009
SHNIH022
SHNIH038
SHNIH030
SHNIH048
SHNIH036
SHNIH034
SHNIH055
SHNIH039
SHNIH051
SHNIH042
SHNIH023
SHNIH045
SHNIH043
SHNIH002
SHNIH053
SHNIH003
Clade A
Clade B

**Figure S8**. **Phylogenetic tree of all 126 staphylococcal genomes based on the polymorphic sites detected in sequence alignment of select core genes (N = 665).** Bootstrap values of at least 80 percent are shown with red dots. For each species-level phylogenetic tree shown in Figure 5, the other two species served as an outgroup for rooting the tree based on this genus-level tree. Body site, healthy volunteer, and species of each genome are shown as sidebars. Refer to Fig. S1 for body site details.

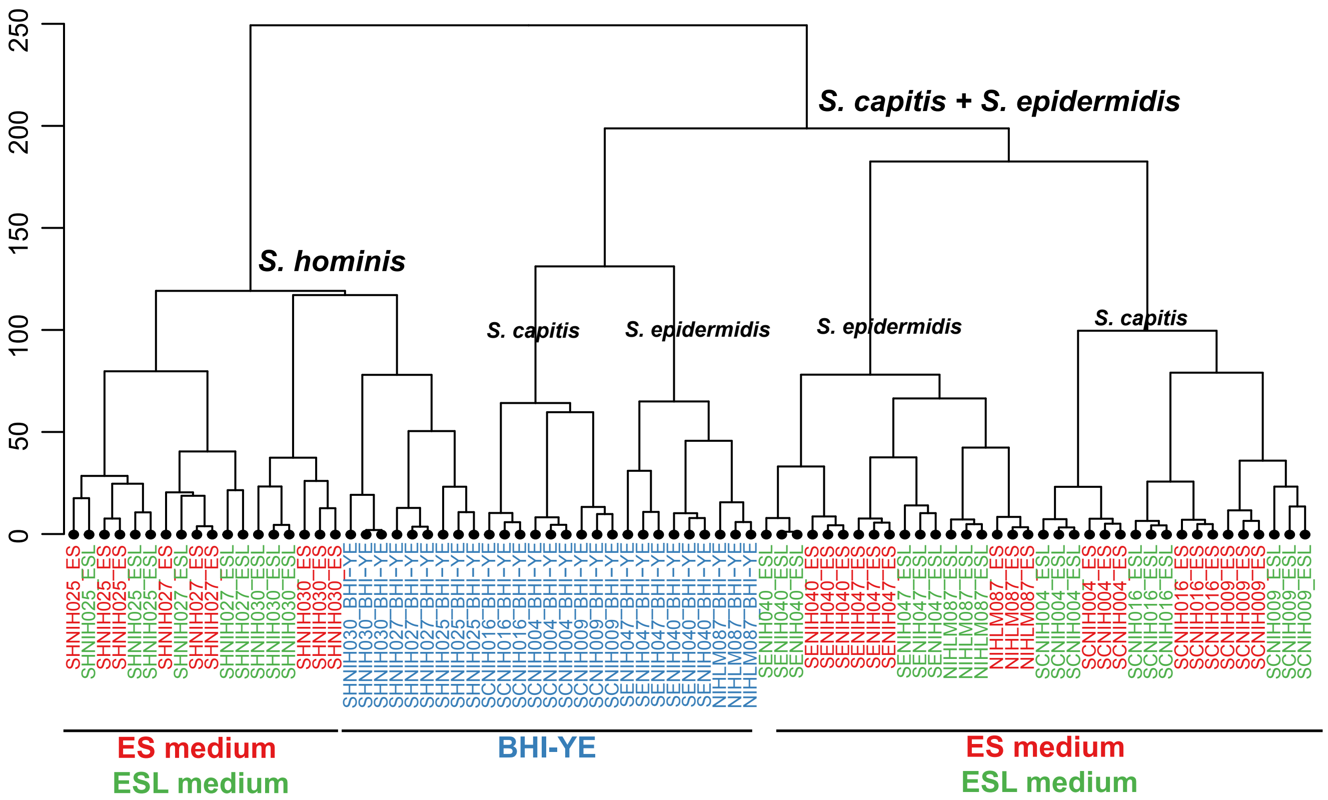

0
50
100
150
200
250
S. capitis + S. epidermidis
S. hominis
S. capitis
S. epidermidis
S. epidermidis
S. capitis
ES medium
ESL medium
BHI-YE
ES medium
ESL medium

**Figure S9 Hierarchal clustering of variance stabilizing transformation-normalized reads for 1647 genus-core genes from all RNA-seq samples (Euclidean distance; Ward)**.

The x-axis represents sample clusters, and the y-axis represents the distances between samples. Colors depict different growth media. Red: ES, Green: ESL, and Blue is BHI-YE. Samples clustered by species (shown as labels) and by growth medium.

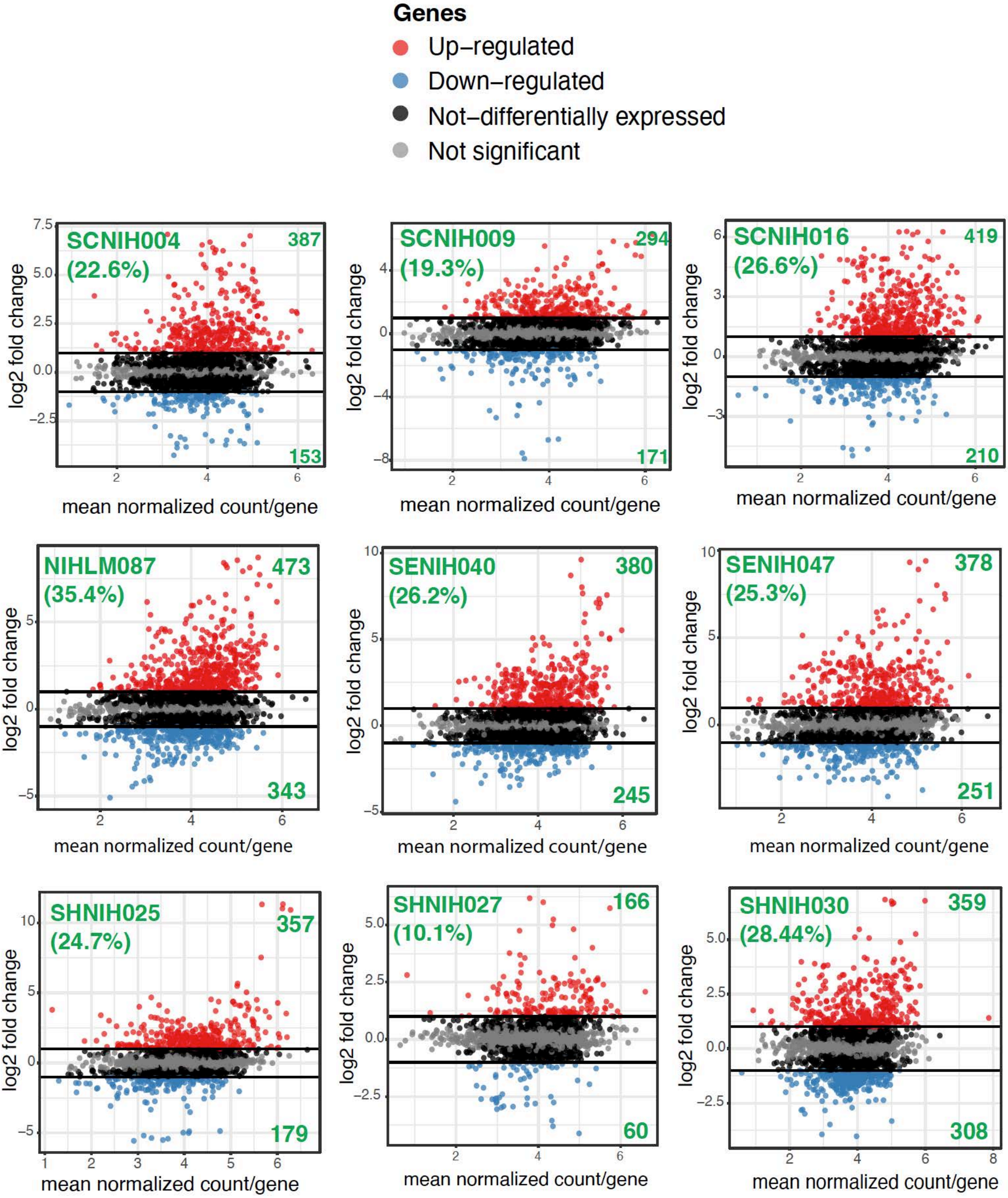

Genes
Up−regulated
Down−regulated
Not−differentially expressed
Not significant
SCNIH004 (22.6%)
387
153
SCNIH009 (19.3%)
294
171
SCNIH016 (26.6%)
419
210
NIHLM087 (35.4%)
473
343
SENIH040 (26.2%)
380
245
SENIH047 (25.3%)
378
251
SHNIH025 (24.7%)
357
179
SHNIH027 (10.1%)
166
60
SHNIH030 (28.44%)
359
308
log2 fold change
mean normalized count/gene

**Figure S10 Differential gene expression plot for each isolate.**

Each box represents the fold change in expression of each gene in the ES medium relative to BHI-YE for each isolate plotted against the mean normalized count per gene. Red dots represent upregulated genes, blue represent downregulated genes ($\geq$ or $\leq$ 2-fold change, adjusted P value < 0.05, DESeq). The actual number of upregulated (top) and downregulated (bottom) genes is show in green. The numbers shown in green inside brackets placed below each isolate label represent the percentage of genes within a genome that were differentially regulated in the ES medium relative to BHI-YE.

**Supplementary Table 1 – Prevalence and mean relative abundance of skin-resident staphylococcal species based on 16S rRNA amplicon analysis.**

| Species | Mean Percent Relative Abundance* ± Standard Deviation | Prevalence by subjects (n = 22) (%) | Prevalence by samples (n = 298) (%) |
|---|---|---|---|
| *S. epidermidis* | 52.25 ± 1.87 | 22 (100) | 284 (95.30) |
| *S. capitis* | 26.37 ± 1.77 | 22 (100) | 225 (75.50) |
| *S. hominis* | 23.65 ± 1.82 | 22 (100) | 208 (69.80) |
| *S. warneri* | 8.27 ± 1.23 | 22 (100) | 121 (40.60) |
| *S. lugdunensis* | 3.98 ± 1.69 | 17 (77) | 31 (10.40) |
| *S. haemolyticus* | 6.27 ± 1.49 | 16 (73) | 53 (17.79) |
| *S. auricularis* | 30.79 ± 7.03 | 13 (59) | 27 (9.06) |
| *S. cohnii* | 12.19 ± 3.25 | 13 (59) | 35 (11.74) |
| *S. pettenkoferi* | 9.19 ± 1.92 | 9 (41) | 19 (6.38) |
| *S. aureus* | 8.44 ± 2.83 | 8 (36) | 17 (5.70) |
| *S. saccahrolyticus* | 16.32 ± 3.47 | 6 (27) | 31 (10.40) |
| *S. epidermidis group* | 16.03 ± 2.88 | 6 (27) | 27 (9.06) |
| *S. saprophyticus* | 0.84 ± 0.28 | 6 (27) | 8 (2.68) |
| *S. simulans* | 3.31 ± 1.14 | 5 (23) | 6 (2.01) |
| *S. pasteuri* | 3.26 ± 1.06 | 5 (23) | 14 (4.70) |
| *S. haemolyticus group* | 4.48 ± 1.35 | 4 (18) | 13 (4.36) |
| *S. petrasi* | 0.86 ± 0.22 | 4 (18) | 7 (2.35) |

*Mean percent relative abundance of each species was calculated using only those samples that were positive for the given species (range: minimum 3 reads to maximum 12119 reads).

**Supplementary Table 5 – Genus-core genes with predicted role in skin colonization**

| Gene ID | Genes | Function | Role in skin colonization |
|---|---|---|---|
| S0510 | *srtA* | covalent anchoring of adhesins to the bacterial cell wall | bacterial adhesion, biofilm formation, and immune escape (1) |
| S1463, S0384, S0385, S1644 | *dltABCD* | D-alanylation of teichoic acids | Resistance to host antimicrobial peptides (2) |
| S0319 | *mprF* | phospholipid lysylation | Resistance to host antimicrobial peptides (3) |
| S0957 | *sepA* | protease | AMP degradation (4) |
| S1750, S1576 | *vraF, vraG* | AMP export | Resistance to host antimicrobial peptides (5) |
| S1493, S1721, S1590 | *graR, graS, graX* | Aps system | AMP sensor, regulator of AMP resistance mechanisms (6) |
| S0518- S0520 | *capABC* | Poly-γ-DL-glutamate capsule biosynthesis | Protects from AMPs, phagocytosis, and high salt concentration (7) |
| S1387 | *oatA* | O-acetylates peptidoglycan | Lysozyme resistance (8) |
| S0663 | *vraX* | binds host complement protein | Inhibit classical complement pathway (9) |
| S1611 | *atlE* | Bifunctional autolysin/adhesin | biofilm formation, vitronectin binding (10) |
| S1620, S1526, S1030, S1031 | *pmtABCD* | ABC transporter for all Phenol-soluble modulins (PSM) classes | Virulence, biofilm (11) |
| S0455, S1812 | - | Phenol-soluble modulins-beta | promote biofilm maturation and dissemination (12) |

| | | | |
|---|---|---|---|
| S0624, S0625, S1613, S1766, S0626, S0627, S1813 | *tagDXBGHA* | poly-glycerol-phosphate teichoic acids synthesis | Role in nasal colonization (13) |
| S1337 | *gehC* | lipase | establish residence in the hair follicles (14) |
| S0557 | *betA* | Choline dehydrogenase | biosynthesis of the osmoprotectant glycine betaine (15) |
| S0487, S0486, S0485, S1232 | *opuCABCD* | Glycine betaine/choline/carnitine transport | Osmoprotolerance (16) |
| S1313, S0867-S0872<br>S0873 | *ureABCEFGD*<br><br>*yut* | urease-production<br><br>urea transport | acid response and pH homeostasis (17) |
| S0947- S0949 | *yfmCDE* | ferric citrate transporter | Iron acquisition (18) |
| S0951 | *isdG* | liberates iron from host heme | Iron acquisition (19) |

**Supplementary Table 8. Composition of artificial skin media** (www.pickeringlabs.com/)

1. **Eccrine Sweat (ES):**

Artificial Eccrine Perspiration (pH 5.5); Catalog Number: 1700-0023

**Amino Acids**
Concentrations for listed amino acids range from 0.002 g/L (for Taurine) to 0.30 g/L (for Serine)

- Glycine
- L-Alanine
- L-Arginine
- L-Asparagine
- L-Aspartic acid
- L-Citrulline
- L-Glutamic acid
- L-Histidine
- L-Isoleucine
- L-Leucine
- L-Lysine as hydrochloride
- L-Methionine
- L-Ornithine as hydrochloride
- L-Phenylalanine
- L-Serine (Largest amount)
- L-Threonine
- L-Tyrosine
- L-Valine
- Taurine

**Metabolites**
Concentration for listed metabolites range from 0.015 g/L (for Uric Acid) to 1.74 g/L (for Urea)

- Uric acid
- Urea
- Lactic acid
- Ammonia

**Minerals**

- Sodium – 33 mmole/L
- Zinc – 11.21 µmole/L
- Chloride – 80.34 mmole/L
- Calcium – 5.49 mmole/L
- Iron – 4.62 µmole/L
- Magnesium – 1.67 mmole/L
- Potassium – 33 mmole/L
- Sulfate – 2.57 mmole/L

2. **Eccrine Sweat with lipids (ESL):** This medium was prepared using ES medium as the base. Tween 80 was added at 0.1%. The following sebum/apocrine emulsion was added at 1% for final growth curves.

**Apocrine sweat**: emulsion of sebum plus other ingredients. Catalog Number 1700-0556

| Compound | Concentration (g/L) |
|---|---|
| L-Alanine | 0.1-0.5 |
| L-Aspartic acid | 0.01-0.1 |
| L-Citrulline | 0.1-0.5 |
| L-Glutamic acid | 0.5 -2 |
| L-Glutamine | 0.1-0.5 |
| Glycine | 0.1-0.5 |
| L-isoleucine | 0.1-0.5 |
| L-Leucine | 0.1-0.5 |
| L-Lysine | 0.5-2 |
| L-Phenylalanine | 0.01-0.1 |
| L-Proline | 0.1-0.5 |
| L-Serine | 0.1-0.5 |
| L-Threonine | 0.1-0.5 |
| L-Tryptophan | 0.1-0.5 |
| L-Tyrosine | 0.01-0.1 |
| L-Valine | 0.1-0.5 |
| Creatine | 0.01-0.1 |
| Urea | 0.2-2 |
| Citric acid | 0.1-0.5 |
| Formic Acid | 0.01-0.1 |
| Lactic Acid | 0.5 - 2 |
| Glucose | 0.01-0.1 |
| Butyric acid | 2-3 |
| Valeric acid | 2-3 |
| a-hydroxy-n-butyric acid-sodium salt | 0.01-0.1 |
| 3-hydroxybutyric acid | 0.01-0.1 |
| a-hydroxy-iso-butyric acid | 0.1-0.5 |
| $(NH_4)_2SO_4$ | 0.1-0.5 |
| $CaCl_2$*$2H_2O$ | 0.5 - 2 |
| $CuSO_4$*$5H_2O$ | 0.001-0.01 |
| $Fe(NO_3)_3$*$9H_2O$ | 0.001-0.01 |
| $MgCl_2$*$6H_2O$ | 0.1-0.5 |
| NaCl | 0.5 - 2 |
| $ZnCl_2$ | 0.001-0.01 |

**Sebum components**

- Palmitic acid (CAS# 57-10-3): 0.5 % w/v

- Stearic acid (CAS# 57-11-4): 0.25% w/v
- Oleic Acid (CAS# 112-80-1): 0.9 % w/v
- Linoleic acid (CAS# 60-33-3): 0.25% w/v
- Coconut oil (CAS# 8001-31-8): 0.75 % w/v
- Olive oil (CAS# 800-25-0): 1% w/v
- Paraffin Wax (CAS# 8002-74-2): 0.5 % w/v
- Synthetic Spermacetti: 0.75% w/v
- Squalene (CAS# 111-02-4): 0.25% w/v
- Cholesterol (CAS# 57-88-5): 0.25% w/v
- Triethanolamine (CAS#102-71-6): 0.8% w/v

**Supplementary Table 9. Distribution of differentially expressed genes in ES medium relative to BHI-YE between core and accessory gene partitions**

Differentially expressed genes = Fold change ≥ or ≤ 2, adjusted P value < 0.05

| Species | Isolate | No. of genus-core genes | Percent (%) | No. of species-restricted core genes | Percent (%) | No. of accessory genes | Percent (%) |
|---|---|---|---|---|---|---|---|
| *S. epidermidis* | NIHLM087 | 583 | 72 | 149 | 18 | 81 | 10 |
| | SENIH040 | 428 | 70 | 107 | 17 | 78 | 13 |
| | SENIH047 | 398 | 64 | 119 | 19 | 104 | 17 |
| *S. capitis* | SCNIH004 | 345 | 66 | 149 | 29 | 28 | 5 |
| | SCNIH009 | 268 | 58 | 131 | 29 | 62 | 13 |
| | SCNIH016 | 438 | 70 | 172 | 27 | 18 | 3 |
| *S. hominis* | SHNIH025 | 371 | 70 | 70 | 13 | 92 | 17 |
| | SHNIH027 | 166 | 74 | 32 | 14 | 26 | 12 |
| | SHNIH030 | 463 | 70 | 85 | 13 | 115 | 17 |

**Datasets in Excel format**

**Supplementary Table 2.** Genome statistics of isolates used in the current study

**Supplementary Table 3.** Gene presence absence matrix based on individual species pan-genome

**Supplementary Table 4.** Gene presence absence matrix of merged genus pan-genome

**Supplementary Table 6.** List of species-restricted core genes detected in genus pan-genome.

**Supplementary Table 7.** pan-GWAS analysis showing clade-specific gene enrichment in each species

**Supplementary Table 10.** List of differentially expressed genes in the ES medium relative to BHI-YE per isolate
